# Enhanced instructed fear learning in delusion-proneness

**DOI:** 10.1101/264739

**Authors:** Anaïs Louzolo, Rita Almeida, Marc Guitart-Masip, Malin Björnsdotter, Martin Ingvar, Andreas Olsson, Predrag Petrovic

**Affiliations:** Department of Clinical Neuroscience, Karolinska Institutet, Stockholm, Sweden; Department of Neuroscience, Karolinska Institutet, Stockholm, Sweden; Department of Neurobiology, Care Science and Society, Karolinska Institutet, Stockholm, Sweden

**Keywords:** delusion-proneness, predictive coding, instructed conditioning, top-down learning, expectation bias, social learning, fMRI

## Abstract

Psychosis is characterized by distorted perceptions and deficient low-level learning, including reward learning and fear conditioning. This has been interpreted as reflecting imprecise priors in a predictive coding system. However, this idea is not compatible with formation of overly strong beliefs and delusions in psychosis-associated states. A reconciliation of these paradoxical observations is that these individuals actively develop and use higher-order beliefs in order to interpret a chaotic environment. In the present behavioural and fMRI study, we compared delusion-prone individuals (n=20), a trait related to psychotic disorders, with controls (n=23; n=20 in fMRI-part) to study the effect of beliefs on fear learning. We show that instructed fear learning, involving explicit change of beliefs and an associated activation of lateral orbitofrontal cortex, is expressed to a higher degree in delusion-prone subjects. Our results suggest that strong high-level top-down learning co-exists with previously reported weak low-level bottom-up learning in psychosis-associated states.

## Introduction

Clinical observations of patients with psychosis suggest that these individuals have difficulties to focus on one stimulus at a time, especially in an acute psychotic state. Instead, their attention often quickly shifts between different irrelevant stimuli that they perceive as highly salient. The same individual may simultaneously have a set of delusions that are resistant to change, despite being extremely unlikely or even bizarre to most people. The paradox that poorly reliable low-level processes (such as unstable perceptions) co-exist with overly stable high-level beliefs (such as delusions) is of a central question in psychosis research (Schmack et al. 2013). Here, we used a combined instructed fear learning (Mertens et al. 2016; Phelps et al. 2001) and classical fear conditioning (Fullana et al. 2016) task to test whether belief formation is stronger in delusion-proneness, a trait associated to psychotic disorders that is expressed in healthy subjects (van Os et al. 2009; Peters et al. 2004), than in controls.

Mirroring the clinical picture of unstable perceptions, experimental research supports the idea that low-level processes are dysfunctional in schizophrenia and related endophenotypes (Javitt & Freedman 2015). A consequence of noisy perceptual processes would be a less efficient bottom-up learning. This has been suggested for psychosis-related states in various simple learning paradigms including associative learning (Corlett et al. 2007; Corlett & Fletcher 2012), reward learning (Murray et al. 2008; Roiser et al. 2009; Schlagenhauf et al. 2014) and fear conditioning (Balog et al. 2013; Holt et al. 2009; Holt et al. 2012; Jensen et al. 2008; Romaniuk et al. 2010). These studies on patients and related endophenotypes have often shown both a smaller learning effect of the true association and an increased learning effect of non-existent associations, in line with the aberrant salience hypothesis (Kapur 2003).

Cognitive neuroscience research on distorted perceptions and deficient low-level learning related to psychosis has recently focused on the involvement of expectations (or priors) in underlying mechanisms (Adams et al. 2013; Fletcher & Frith 2009). It has been suggested that expectations are fundamental for interpreting input from the external and internal environments that are often noisy or incomplete (Friston 2005). Whenever expectations and incoming signals do not match, an error signal is generated that promotes an adjustment of expectations or input processing until the error is minimized. This will theoretically lead to the most optimal representation of the world at a given time. The *hierarchical predictive coding hypothesis* suggests that comparisons between input signals and expectations occur at all levels of the brain networks, including low- and high-level networks (Friston 2005). Error signals from low-level processes propagate in this hierarchy until higher-level priors can account for them. Apart from being essential for normal information processing and any type of learning, it has been proposed that this organization may mechanistically explain psychotic symptoms (Adams et al. 2013; Fletcher & Frith 2009). Specifically, it has been suggested that the balance between bottom-up signals and top-down influence of expectations is altered in psychotic states (Adams et al. 2013; Fletcher & Frith 2009) due to aberrant (or hyper) salience of incoming information (Kapur 2003) - possibly linked to a hypersensitive dopamine system (Kuepper et al. 2012) - and weakened or imprecise priors (Adams et al. 2013; Fletcher & Frith 2009). Recently, hierarchical Bayesian models have been successfully applied to explain hallucinations and underlying processes observed in psychosis-associated states (Powers et al. 2017). Importantly, predictive coding models have so far not been able to account for both chaotic perceptions (involving imprecise priors) and delusions (involving overly precise priors).

In contrast to bottom-up learning, recent studies suggest that the effect of high level top-down learning is stronger in patients with psychosis and delusion-prone subjects than in healthy controls (Schmack et al. 2013; Teufel et al. 2015). Namely, after being presented with higher order information, these phenotypes use high-level priors in a top-down fashion more readily than controls, in order to interpret simple perceptual input (Schmack et al. 2013; Teufel et al. 2015). We propose that the preponderance to integrate higher order information may also lead to stronger explicit belief formation, and ultimately to delusions. Moreover, it has been suggested that the formation of overly strong beliefs and delusions is a secondary consequence of adaption to aberrant low-level signals (Kapur 2003). We suggest that a strategy of integrating explicit information in a proactive manner to facilitate interpretation of a noisy environment, may also be important for belief formation in psychosis-related states.

Here, we tested the effect of prior explicit information manipulation (involving a change of conscious expectations) on *social fear learning* in subjects with high delusion-proneness and matched controls. We hypothesized that explicitly induced expectations about the threat value of specific social stimuli would have stronger effect on affective learning in delusion-prone participants, in sharp contrast to previously performed fear conditioning studies on psychosis patients (Holt et al. 2009; Holt et al. 2012; Jensen et al. 2008; Romaniuk et al. 2010), and schizotypal individuals (Balog et al. 2013), which have suggested a weaker learning. Our main measure consists of explicit evaluation of social stimuli, and involves, therefore, conscious beliefs about the context. In line with previous studies where a change in expectations underlies a change in emotional experience (Eippert et al. 2007; Golkar et al. 2012; Kanske et al. 2011; Wager et al. 2008), we hypothesized that these effects would be related to prefrontal, in particular lateral orbitofrontal cortex, as well as its interaction with regions processing pain and fear.

### The current study

The present behavioural and functional brain imaging study combined both instructed fear learning (top-down learning) (Mertens et al. 2016; Phelps et al. 2001) and classical fear conditioning (bottom-up learning) (Fullana et al. 2016). We used four pictures of neutral male faces as our conditioned stimuli (CS); two would be paired with an aversive unconditioned stimulus (UCS) (i.e. CS+) and two would not be reinforced (i.e. CS-). In the *Instruction phase*, information regarding UCS contingencies was presented for two of the CS:s (iCS+ and iCS-). No information was provided for the other two other CS:s (niCS+ and niCS-). In the *Acquisition phase*, both CS+ were paired with the UCS with a 50% reinforcement rate. Finally, in the *Extinction phase* all CS were presented without any UCS pairing. Our main behavioural outcome variable was evaluative likability ratings of the CS:s before and after each phase. The difference score between CS- and CS+ (for instructed and non-Instructed CS-pair) after each phase is referred to as the *affective learning index*. Apart from the ratings, we also measured the skin conductance response (SCR), serving as a physiological index of affective learning. Finally, we analysed the underling brain activations using functional magnetic resonance imaging (fMRI) for the Acquisition phase. See the Method section and Fig 1 for more detailed information.

**Fig. 1.**
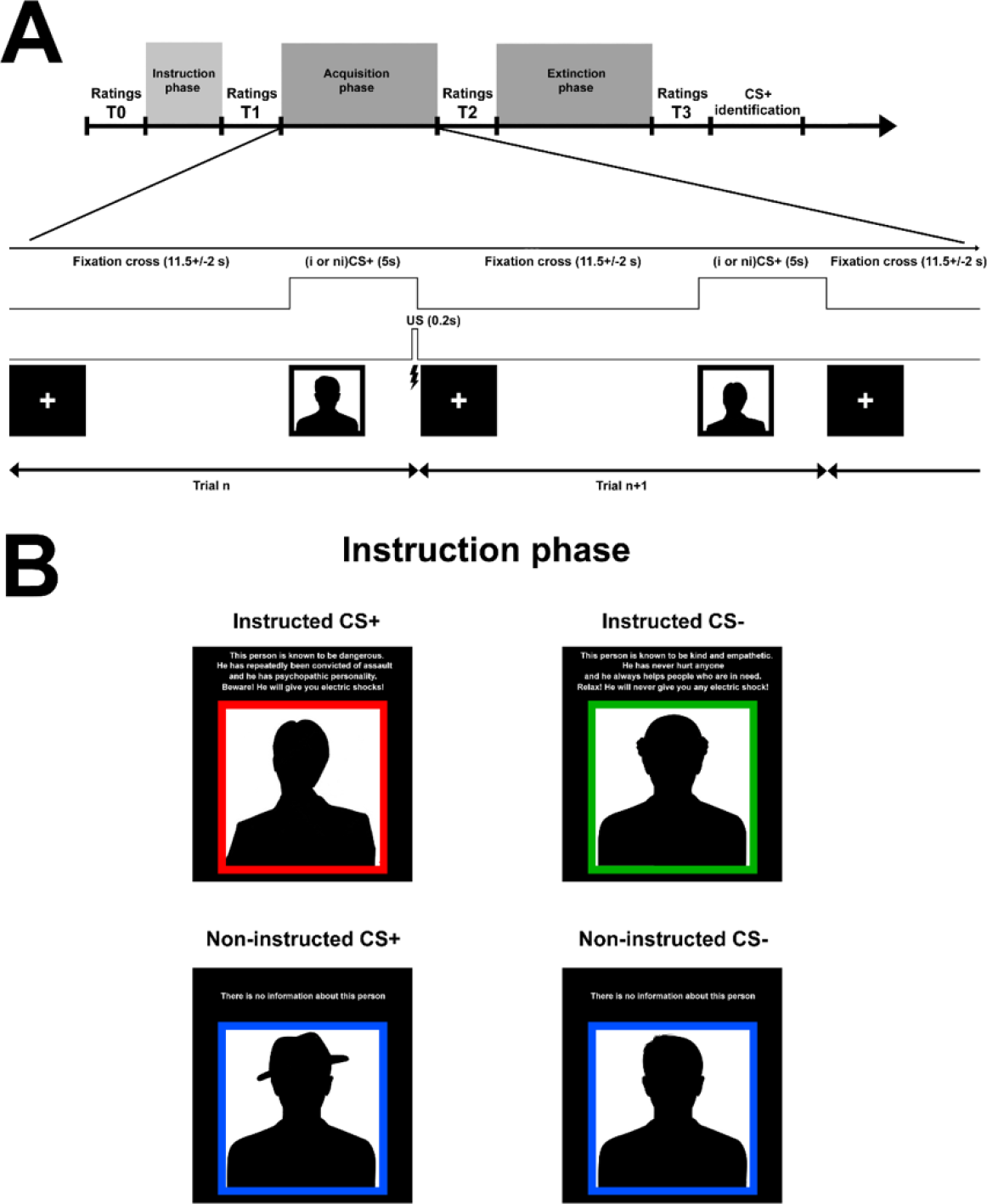
Subjects and experimental design. **(A)** Timeline of paradigm. The *acquisition and extinction phases* each CS was displayed 12 times for 5 seconds, and the jittered inter-trial interval was 11.5± 2 seconds. The CS+ were coupled with UCS (mildly painful electric stimulation) with a 50% contingency in the acquisition phase and there was no UCS in the extinction phase. Participants were asked to rate how friendly each CS was experienced, using a visual analogue scale (−100 to 100; Methods). In order to estimate learning in our paradigm we calculated the difference between CS- rating and CS+ rating for each CS-pair (instructed and non-instructed). This difference score is referred to as “*affective learning index*” and the main outcome value in the study. We analysed four *affective learning indices*: 1) T0: before instruction learning 2) T1: after instruction learning, 3) T2: after acquisition and 4) T3: after extinction. All ratings were normalized in regards to T0. **(B)** In the *instruction phase*, two of the faces (instructed CS+ and CS-; iCS+/iCS-) were coupled with information about their contingencies with the UCS that included a fabricated short description about their personality and the risk of being associated with an aversive stimulation. The two other CS faces (non-instructed CS+ and CS-; niCS+/niCS-) contained no information about their contingencies with the UCS. Instructions were presented twice (followed by ratings – T1’ and T1) in

## Results

### 1. Behavioural results

#### Ratings

##### Baseline ratings

A baseline rating (T0) was collected for each face before any information was presented and it was used for normalisation of subsequent ratings (Fig 1A). We tested whether groups (control group and delusion-prone group) differed on the averaged absolute value of the initial ratings, and found no significant difference (t=0.092, p=0.927, independent two-sample t-test) (Fig S1). This result suggests that group differences associated to instructions or conditioning cannot be explained simply by a difference between the groups in their general rating strategy.

##### All phases together

We tested our main hypothesis, i.e. that delusion-prone participants would show a larger effect of instructions compared to controls, by performing a repeated-measure linear model (phases x group) on the *affective learning index* for the instructed stimuli, between the three phases (T1, T2 and T3). In line with our prediction we found a significant group effect (t=− 2.34, df=46.83, p=0.012, one-tailed) indicating that the overall effect of instructions was significantly larger in delusion-prone individuals (mean=125.77, SD=93.06) than in the control group (mean=74.50, SD=67.98) (Fig. 2A and B), but no significant main effect of phases, nor any significant phases by group interaction.

**Fig. 2.**
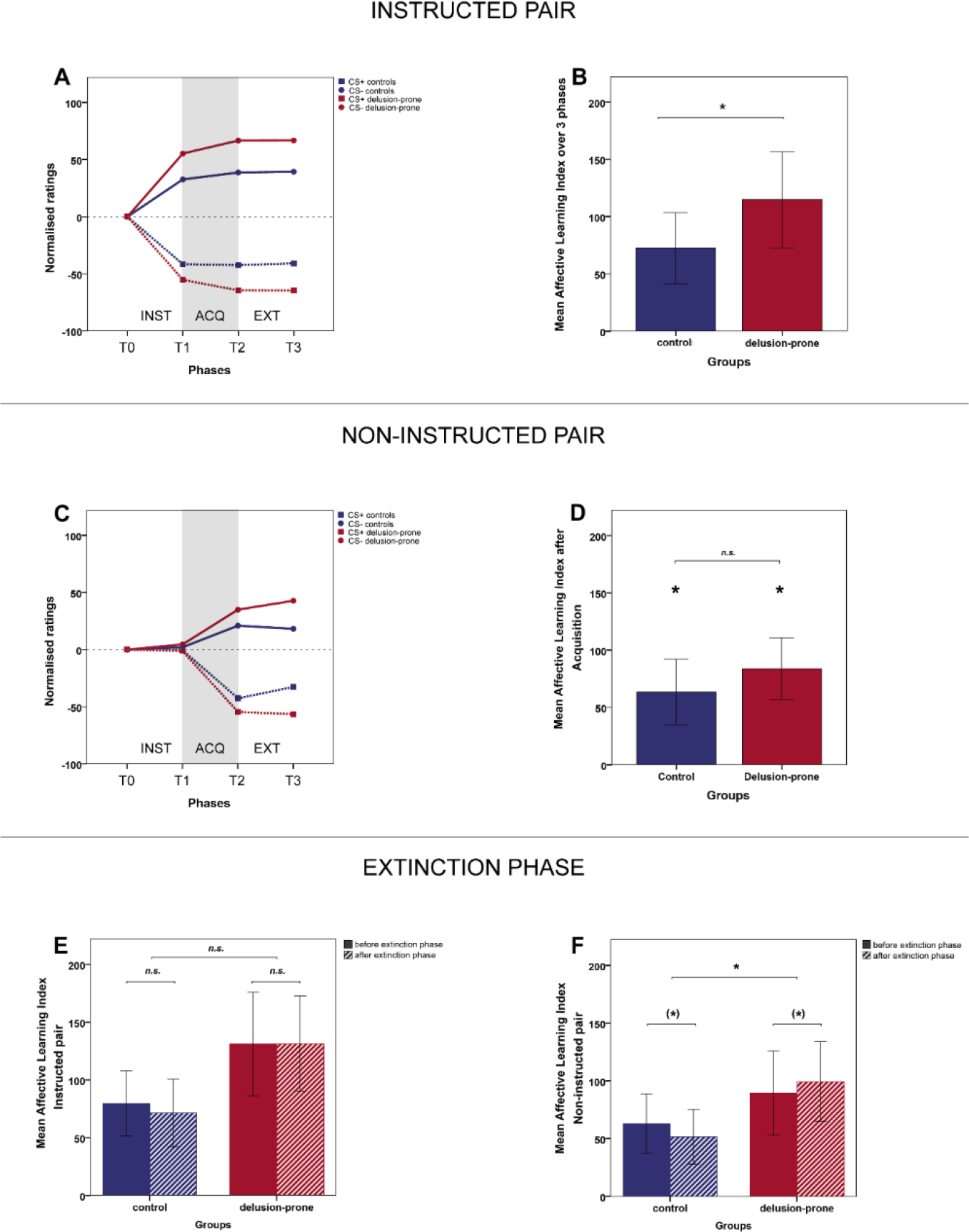
Behavioural results of instructed and non-instructed learning. **(A)** Timeline of likability ratings for instructed CS-stimuli over the three phases. **(B)** Instructed *affective learning index* average over the three phases in controls (mean=74.50, SD=67.98), and delusion-prone individuals (mean=125.77, SD=93.06). In line with our prediction we found a significant group effect (t=-2.34, df=46.83, p=0.012, one-tailed) indicating that the overall effect of instructions was significantly larger in delusion-prone individuals. **(C)** Timeline of likability ratings for non-instructed CS-stimuli over the three phrases. **(D)** A repeated-measure linear model on the non-instructed *affective learning index*, between groups, before and after the acquisition phase (T1 and T2) showed no group effect (t=-1.46, df=71.03, p=0.148) although there was a general effect of acquisition (t=-5.742, df=41, p<0.001). The average non-instructed *affective learning index* after acquisition was somewhat larger (albeit non-significant) in the delusion-prone group than in the control group (DG: mean=89.45, SD=81.52; CG: mean=63.00, SD=62.16). The *Affective learning index* for the non-instructed CS pair increased significantly after acquisition (T2 vs T1) both in controls (mean T1=-0.261, SD=38.46; mean T2=63.00, SD=62.16; paired t-test t=-4.405, df=22, p<0.001 one-tailed) and in delusion-prone individuals (mean T1=5.70, SD=47.92; mean T2 =89.45, SD=81.52; paired t-test t=-6.165, df=19, p<0.001 one-tailed). **(E)** Instructed *affective learning index* before and after the extinction phase (T2 and T3) in delusion-prone (mean T2= 131.15, SD=100.35; mean T3= 131.40, SD=92.07; paired t-test t=-0.048, df=19, p=0.96) and controls (mean T2=79.74, SD=67.92; mean T3= 71.26, SD=70.26; paired t-test t=1.04, df=22, p=0.31). No significant group differences was observed. **(F)** Non-instructed *affective learning index* before and after the extinction phase (T2 and T3) in delusion-prone (mean T2= 89.45, SD=81.52; mean T3=99.35, SD=87.00; paired t-test t=-1.78 p=0.09) and controls (mean T2=63.00, SD=62.16; mean T3= 51.65, SD=56.42; paired t-test t=1.63, df=22, p=0.059 one-tailed). A significant interaction between the groups in the extinction effect was observed for the non-instructed stimulus pairs (t=2.339, df=41. p=0.024 - repeated-measure linear model (T2/T3phases x group) on the non-instructed *affective learning index)*. **Error bars: 2 S.E**

##### Instruction phase

Within the instruction phase (Fig 1), instructions were presented twice to increase learning, with a likability rating after each presentation (T1’ and T1). For the remaining of the analyses we only used the second ratings as the instruction phase rating (referred as T1) indicating the total learning effect in this phase. However, when specifically studying the *affective learning index* after the first instruction were given (T1’) we observed a clear learning effect for the instructed stimuli (iCS+/iCS-) both for the control group (mean=50.65, SD=79.06, one-sample t test t=3.073, df=22, p=0.003 one-tailed) and the delusion-prone individuals (mean=97.40, SD=85.89, one-sample t test t=5.07, df=19, p<0.001 one-tailed) (Fig. S2A) suggesting an effect of instructions in both groups for T1’. An independent sample t-test revealed a significant group difference (t=-1.858, df=41 p=0.035 one-tailed) (Fig. S2A). In both groups, *affective learning* increased significantly after the second instruction presentation (T1) (*control group*: mean=72.52, SD=74.59, paired t-test t=-1.963; df=22, p=0.032 one-tailed; *delusion-prone* mean=114.75, SD=93.26, paired t-test t=-2.350, df=19, p=0.015 one-tailed) suggesting an effect of instructions in both groups also for T1. The group difference was on the border of significance (independent t-test t=-1.649, p=0.053 df=41 one-tailed) (Fig. S2B).

##### Acquisition phase

*Affective learning index* for the non-instructed CS pair increased significantly after acquisition (T2 vs T1) in controls (mean T1=-0.261, SD=38.46; mean T2=63.00, SD=62.16; paired t-test t=-4.405, df=22, p<0.001 one-tailed) and in delusion-prone individuals (mean T1=5.70, SD=47.92; mean T2 =89.45, SD=81.52; paired t-test t=-6.165, df=19, p<0.001 one-tailed) (Fig. 2C and D) suggesting an effect of conditioning in both groups. A repeated-measure linear model on the non-instructed *affective learning index*, between groups, before and after the acquisition phase (T1 and T2) showed no group effect (t=-1.46, df=71.03, p=0.148) although there was a general effect of acquisition (t=-5.742, df=41, p<0.001).

A trend towards a conditioning effect based on *affective learning index* was also observed for the instructed CS pair both in controls (mean before acquisition=72.52, SD=74.59; mean after acquisition=79.73, SD=67.93; paired t-test t=-1.679, df=22, p=0.054 one-tailed) and delusion-prone subjects (mean before acquisition=114.75, SD=93.26; mean after acquisition=131.15, SD=100.35 – paired t-test t=-1.704, df=19, p=0.053 one-tailed). A repeated-measure linear model on the instructed *affective learning index*, between groups, before and after the acquisition phase (T1 and T2) showed no group interaction.

The total *affective learning* after conditioning (T2) was larger for the instructed than the non-instructed conditions in the delusion-prone group (mean instructed *affective learning index* =131.15, SD=100.35; mean non-instructed *affective learning index* =89.45, SD=81.52; paired t-test t=2.198, df=19, p=0.041). However, this was not the case in the control group (mean instructed *affective learning index* =79.74, SD=67.93; mean non-instructed *affective learning index* =63.00, SD=62.16; paired t-test t=1.000, df=22, p=0.328) although there was no significant difference of these effects between groups (Fig S3C and D).

##### Extinction phase

We also tested whether *affective learning* was more resistant to extinction after fear conditioning in delusion-prone versus control subjects. A repeated-measure linear model (group x T2/T3-Phase) analysis of *affective learning index* showed a significant interaction between the groups for the non-instructed CS pairs (t=2.339, df=41, p=0.024) suggesting a relatively smaller effect of extinction in the delusion-prone group. Interestingly, while the control group showed a trend towards an expected extinction effect (t=1.63, df=22, p=0.059 one-tailed paired t-test), the delusion-prone group tended to show an opposite effect, i.e. increased *affective learning index* after extinction (t=-1.78 p=0.09) (Fig. 2F).

A similar repeated-measure linear model (group x T2/T3-Phase) analysis of *affective learning index* for the instructed CS pair did not reveal any significant phase by groups interaction (i.e. extinction). The paired t-tests performed on the instructed *affective learning index* after acquisition (T2) and after extinction (T3) in each group did not reveal any significant difference (delusion-prone group: t=-0.048, df=19, p=0.96, paired t-test; control group: t= 1.04, df=22, p=0.31, paired t-test) (Fig. 2E). Thus, both groups showed resistance to extinction for the instructed CS pair.

#### Skin conductance

A one-tailed t-test on the differential SCR (SCR-CS+ vs SCR-CS-) in the acquisition phase for all subjects together, was significantly different from zero (average=0.0151, SD=0.0271; t=3.424, df=37, p=0.001 one-tailed) suggesting a significant conditioning. This was also the case for each group, when analysed separately (controls mean=0.0126μS, SD=0.0248, one-sample t-test t=2.145, df=17, p=0.024 one-tailed - delusion-prone mean=0.0174μS, SD=0.0296, one-sample t-test t=2.628, df=19, p=0.009 one-tailed) (Fig. S3A). There was no group difference (independent two-sample t-test t=-0.741, df=73, p=0.461) (Fig. S3A).

The differential SCR was mainly driven by the iCS-pair as suggested by a significant difference between the instructed and non-instructed condition in controls (instructed mean= 0.0266μS, SD=0.036, non-instructed mean=-0.015μS, SD=0.029; paired t-test t=2.780, df=17, p=0.014) and in delusion-prone individuals (instructed mean= 0.0251μS, SD=0.031, non-instructed mean=0.010μS, SD=0.036; paired t-test t=2.188, df=19, p=0.042). However, there was no significant interaction between the groups (Fig. S3B).

During the extinction phase the differential SCR was no longer significantly different from zero for all subjects together (one-sample t-test t=-1.115, df=75, p=0.268).

Overall, it should be noted that the SCR data recorded in the fMRI scanner was noisy. We only used participants who showed a SCR to at least 20% of the presentations of each CS (hence, considered as responders; n=38). However, many of them were characterised by a low reactivity.

#### Effects of PDI sub-scores on ratings

In an exploratory analysis, we investigated whether PDI scores and their components (distress, preoccupation and conviction) were related to the different ratings for instructed stimuli in the control and delusion-prone group, respectively. For each PDI item that is endorsed, three dimensions are rated by the participant on a 5-point Likert scale (1-5) in order to assess the level of conviction, distress, and preoccupation related to the given item (i.e. conviction, distress, and preoccupation scores, respectively).

In the delusion-prone group we observed a significant correlation between distress scores and the overall instructed *affective learning index* (r=0.555, p=0.011 Pearson correlation tests) (Fig. 3A), as well as the instructed *affective learning index* in each of the three phases: T1 (after instructions): r=0.614, p=0.004; T2 (after acquisition): r=0.518, p=0.019; T3 (after extinction): r=0.571, p=0.009, Pearson correlation tests. While similar correlations were observed for preoccupation and conviction scores they did not reach significance (Fig. S4).

**Fig. 3.**
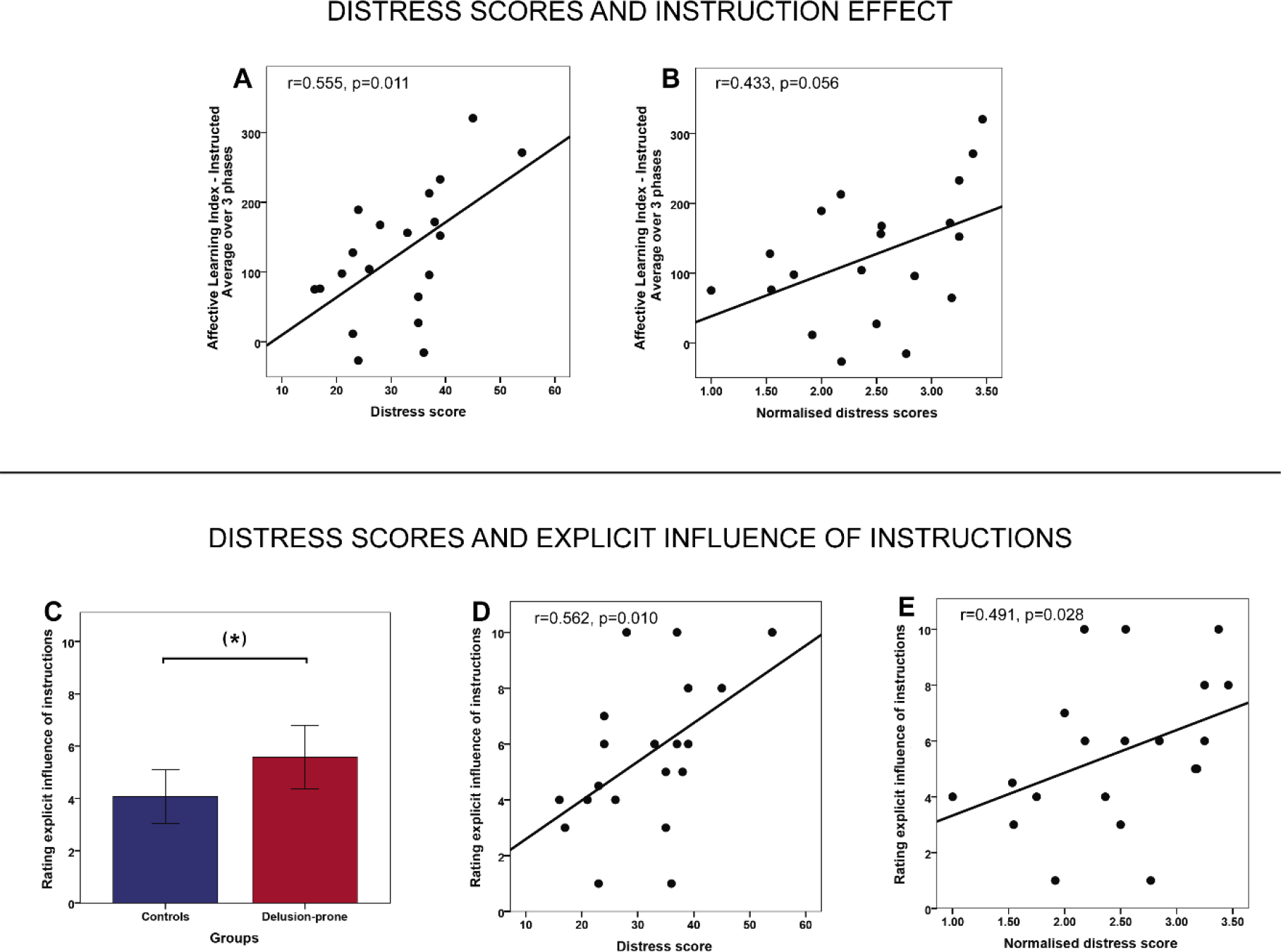
Relation between instruction effects and delusional distress. **(A)** Correlations between *distress scores* and overall instructed *affective learning index* (averaged over three phases) (r=0.555, p=0.011, Pearson correlation tests). **(B)** Correlations between normalised *distress scores* and overall instructed *affective learning index* (averaged over three phases) (r=0.433, p=0.056, Pearson correlation tests). **(C)** Rating of the explicit influence of instructions in controls and delusion-prone individuals. The group difference is on the border of significance t=-1.910, p=0.063, df=40 (independent two-sample t-test). **(D)** Correlation between distress scores and explicit rating of instruction influence in the delusion-prone group (r=0.562, p=0.010, Pearson correlation tests). **(E)** Correlation between normalised distress scores and explicit rating of instruction influence in the delusion-prone group (r=0.491, p=0.028, Pearson correlation tests). **Error bars: 2 S.E**

Since distress seemed as an important variable in relation to effects of instructions in our fear learning paradigm we explored it further. Only analysing the total sum of each of these sub-scores without taking the Yes/No score can be somewhat misleading, as it makes it difficult to differentiate between people who would score high on distress because they have a few delusion-like experiences that are extremely distressing, from people who score as high on distress because they have many delusion-like experiences that are not distressing at all. Normalising to the number of endorsed items (number of “yes” answers, or the so-called “total PDI score”) provides a better estimate of how distressed, preoccupied and convinced participants are, unrelated to whether there is one or several delusion-like experiences. We therefore also compared the control and delusion-prone group in terms of normalised subscores and found that the average normalised distress score in delusion-prone individuals was significantly larger than in the control group (delusion-prone=2.47, control=1.95; independent sample t-test t=-2.593, p=0.013, df=41). Moreover, in the delusion-prone group, the normalised distress score also correlated positively with *affective learning index* after the instruction phases (r=0.527, p=0.017, Pearson correlation tests) (Fig. 3B). This correlation only reached a trend level in the acquisition and extinction phases, as well as when considering the three phases together (r=0.400, p=0.080; r=0.438, p=0.053; r=338, p=0.091, respectively - Pearson correlation tests) (Fig. S5). No significant correlations between normalised distress scores and *affective learning index* were found in the control group.

##### Post-experiment ratings

After the experiment, participants were asked to explicitly rate the influence of instructions, and pain stimuli (respectively) from 0 to 10. An independent sample t-test revealed a trend towards a larger influence of instructions reported by delusion-prone individuals, compared to controls (mean control=4.07, SD=2.42, mean delusion-prone=5.58, SD=2.69; t=-1.910, p=0.063, df=40 two-tailed) (Fig. 3C), while there was no group difference in terms of pain influence.

Interestingly, in the delusion-prone group, the explicit rating of instruction influence was also significantly correlated to the distress sub-score (r=0.562, p=0.01 Pearson correlation tests) (Fig. 3D) and with the normalised distress score (r=0.491, p=0.028 Pearson correlation tests) (Fig. 3E).

#### 2. Imaging results

A simultaneous fMRI measurement showed that the main effect of conditioning (i.e. all CS+ vs all CS- in the acquisition phase) led to activations in brain areas that are consistently reported in fear conditioning studies (Fullana et al. 2016). These included anterior insula, caudal anterior cingulate cortex and thalamus bilaterally as well as brainstem (Fig. 4A; Table S1). However, no significant differences were observed between the groups in the regions of interest (ROI) analysis for (CS+ vs CS-).

**Fig 4.**
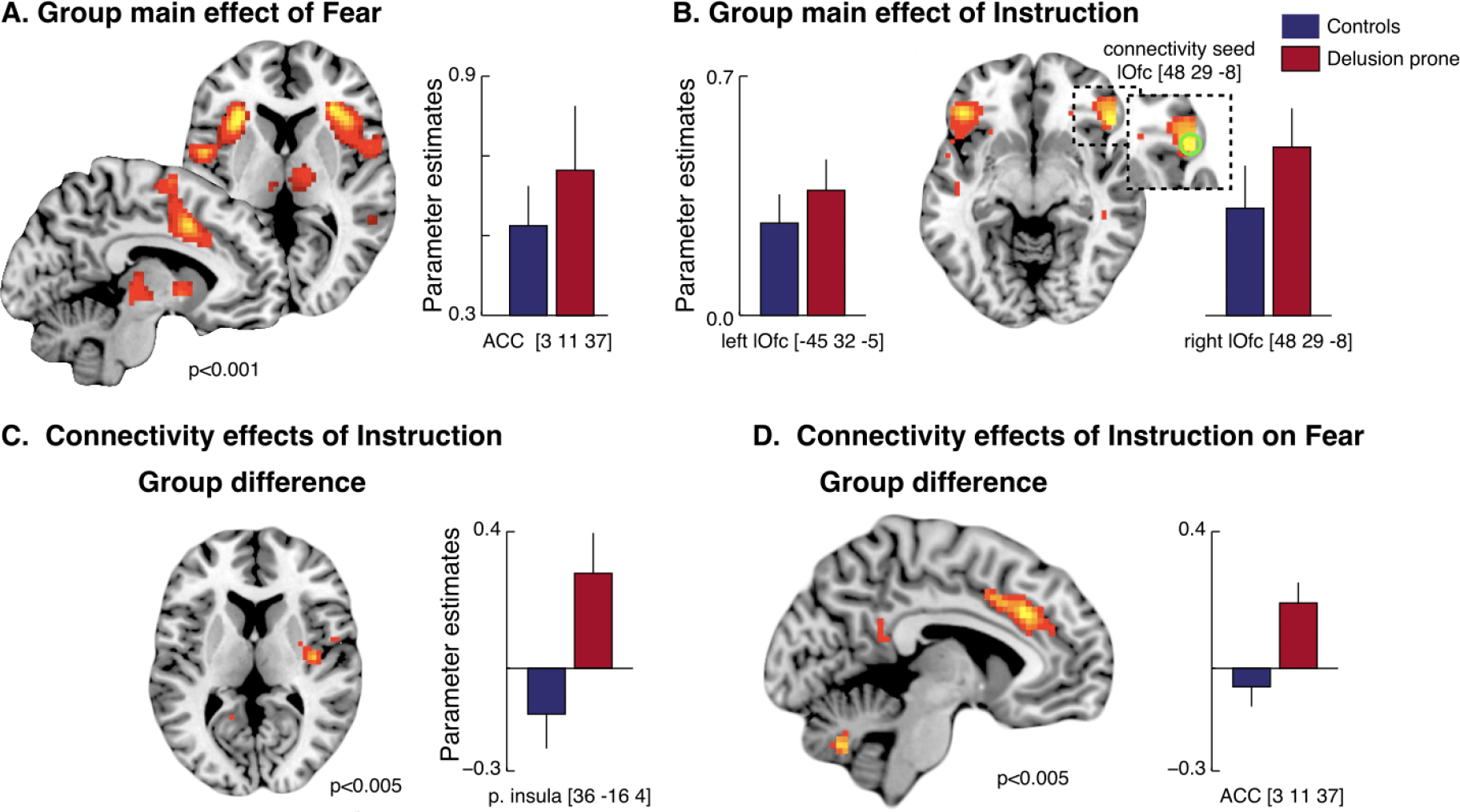
Brain activity related to the effects of conditioning and instructions – BOLD response (A & B) and PPI analyses (C & D). **(A)** Main effect of fear *(CS+ vs. CS-):* an activation in caudal anterior cingulate cortex (cACC), bilateral anterior insula, premotor/dorsolateral prefrontal cortex (dlPFC), right temporoparietal junction (rTPJ) was observed (Table S1). The activation pattern was similar for instructed (iCS+ vs iCS-) and non-instructed (niCS+ vs niCS-) stimuli. No group difference was observed. **(B)** Main effect of instructions: bilateral activations in lateral orbitofrontal cortex (1Ofc) (ROI analysis and whole brain analysis) and an activation in dlPFC (whole brain analysis) were observed (Table S2). This effect was mainly driven by the delusion-prone group. **(C)** A psychophysiological interaction (PPI) analysis on the effect of instructions: an increased connectivity between the right 1Ofc and functionally defined low-level pain processing areas (i.e. right posterior insula) (Z = 3.29, pFWE = 0.004) specifically in delusion-prone individuals was observed. **(D)** A PPI-analysis on the effects of instruction on fear processing: a larger connectivity between the 1Ofc and the cACC (overlapping with fear related activation) in delusion-prone than in control participants (Z = 2.96, pFWE = 0.012) was observed. **Error bars: S.E.**

In line with our hypothesis, we observed a main effect of instructions [(iCS+ + iCS-) vs (niCS+ + niCS-)] in lateral orbitofrontal cortex (lOfc) for all subjects (Fig. 4B; Table S2) - driven mainly by delusion-prone subjects (Fig S6). This suggests a plausible underlying prefrontal mechanism associated with the observed behavioural effects of instructions on fear learning. In addition, delusion-prone individuals also displayed activation in the ventromedial prefrontal cortex (vmPFC) that was not reported in the control group, nor in the all-subject activations (Fig. S6C; Table S2). However, there were no significant differences between the groups in the main effects of instructions (subtraction analysis).

A psychophysiological interaction (PPI) analysis revealed increased connectivity in instructed trials (vs non-instructed trials) specifically for delusion-prone individuals between the right lOfc and functionally defined nociceptive input region (right posterior insula) (Z=3.29, corrected p=0.004), supporting previous findings of an association between sensory processing and lOfc in delusion-prone individuals (Schmack et al. 2013) (Fig 4C). Moreover, PPI-analysis of the effects of instruction on fear processing showed a significantly larger connectivity between the lOfc and the caudal anterior cingulate cortex (cACC), overlapping with fear related activation, in delusion-prone compared to control participants (Z=2.96, corrected p=0.012) (Fig 4D). Last, we tested whether we could replicate the correlation reported in earlier work, between conviction scores and functional connectivity in instructed trials between the right lOfc and functionally defined early sensory processing regions (Schmack et al. 2013) (i.e. right posterior insula, here), specifically for delusion-prone individuals. This analysis showed a significant effect (pFWE=0.003), that was also observed when the PPI-analysis was correlated with the total PDI score (pFWE=0.004) and the normalised convictions scores (pFWE=0.016).

### Discussion

The present findings confirmed our main hypothesis stating that the effect of instructions on fear learning would be larger in delusion-prone individuals than in controls. However, we did not observe any significant group difference in non-instructed fear learning (classical fear conditioning) (Fig. 2B and D). Our results mirror recent studies reporting an increased effect of high-level priors on perceptions in psychosis-related states (Schmack et al. 2013; Teufel et al. 2015) and extend these observations to instructed fear learning (measured with affective ratings). Importantly, as we measured evaluative social ratings we also targeted the participants’ specific beliefs about different social stimuli. Thus, in contrast to the aforementioned studies (Schmack et al. 2013; Teufel et al. 2015) we argue that in psychosis related states, explicit beliefs about the world are also more susceptible to be changed after explicit learning. In addition, we show that delusion-prone individuals displayed a larger *affective learning* than controls, immediately after instructions, i.e. before the CS-UCS pairing. In other words, they had already formed stronger beliefs that biased their experience of the faces, even before low-level learning in the acquisition phase. Thus, we expand previous views on delusion formation as a secondary mechanism in which the individual tries to explain specific aberrant stimuli (Kapur 2003), by suggesting that formation of such beliefs might also represent a pro-active coping strategy in order to facilitate interpretation of an unstable environment.

In the present study we focused on delusion-proneness, a personality trait in healthy individuals that includes subclinical levels of delusional ideation (van Os et al. 2009; Peters et al. 2004). Cognitive, thought- and perceptual mechanisms underlying delusion- and psychosis-proneness are considered to be similar to the one underlying psychosis (Peters et al. 2004; van Os et al. 2009; Teufel et al. 2010; Fusar-Poli et al. 2013). As this phenotype is dimensionally expressed in humans, all individuals are more or less prone to this type of behaviour and related information processing. Thus, this trait has significant impact on variability in human behaviour among healthy subjects. However, similar effects of top-down high-level learning may be present in psychosis patients.

The effect of instructions on fear learning was also significantly related to the degree of delusional distress in the delusion-prone group. This finding was still present when distress scores were normalised, such that they did not depend on the number of endorsed delusional items, which underscores the importance of this dimension in belief formation. These findings may be of special interest since it has been suggested that psychosis-related states characterized with more distress and help seeking are also associated with a larger risk to convert into a clinical psychotic disorder (Fusar-Poli et al. 2013).

The average non-instructed *affective learning index* after acquisition (i.e. evaluative conditioning) was somewhat larger, albeit non-significant, in the delusion-prone group compared to the control group (Fig 2D). At first glance, this result seems to contrast with previous studies showing a smaller classical fear conditioning effect in psychosis patients (Holt et al. 2009; Holt et al. 2012; Jensen et al. 2008; Romaniuk et al. 2010) and schizotypal individuals (Balog et al. 2013). However, it is important to keep in mind that the non-instructed condition may involve a faster development of explicit beliefs about contingencies than in classical fear conditioning due to the presence of an instructed condition in the same experiment. Thus, our non-instructed fear learning cannot be simply compared to classical fear conditioning studies. Future studies will have to control for such confounding effects when comparing instructed versus non-instructed conditions.

We also tested whether *affective learning* was more resistant to extinction in delusion-prone subjects than controls (Fig. 2E and F). Intriguingly, our results showed a significant interaction effect between group and the extinction for the non-instructed CS pair, suggesting that while extinction was present in the controls, the *affective learning index* increased in delusion-prone individuals after the extinction phase. This implies that delusion-prone participants actually reinforce their prior beliefs even when confronted with contradictory evidence. These findings are in line with the *bias against disconfirmatory evidence* described in psychosis-related states, whereby schizophrenia patients (Woodward et al. 2008; Moritz & Woodward 2006; Veckenstedt et al. 2011; Woodward et al. 2006; McLean et al. 2016) and delusion-prone individuals (Buchy et al. 2007; Woodward et al. 2007; Orenes et al. 2012) exhibit a tendency to disregard evidence that goes against the current assumption. Both groups showed a resistance to extinction in the context of instructed stimuli, suggesting that the extinction effect might generally be reduced when part of the fear learning is supported by higher-order beliefs.

Apart from the effects of fear learning measured with *affective learning index*, the subjects also explicitly rated how much the painful stimulation and the instructions affected them. Interestingly, although no group difference was observed for the painful stimulation, delusion-prone subjects tended to rate that they were more affected by the instructions than controls. Also, this effect was significantly correlated with the delusional distress for the instructed stimuli in the delusion-prone group (similarly to the *affective learning index).* Thus, there seems to be a metacognitive awareness in delusion-prone subjects that they are highly affected by explicit information.

Our fMRI results revealed that the main effect of conditioning led to activations in brain areas that are consistently reported in fear conditioning studies including caudal ACC, anterior insula, thalamus and brainstem (Fullana et al. 2016), but no group differences were reported (Fig. 4A; Table S1). In line with our hypothesis, we observed a main effect of instructions in lateral orbitofrontal cortex (lOfc) for all subjects (Fig. 4B; Table S2) - driven mainly by delusion-prone subjects (Fig S6). This suggests a plausible underlying prefrontal mechanism associated with the observed behavioural effects of instructions on fear learning – an effect that was significantly larger in the delusion-prone group than in the control group.

A psychophysiological interaction (PPI) analysis revealed increased functional connectivity in instructed trials specifically for delusion-prone individuals between the right lOfc and functionally defined primary nociceptive input region (right posterior insula), supporting previous findings of an association between sensory processing and lOfc activity in schizophrenia (Schmack et al. 2017) and delusion-proneness (Schmack et al. 2013)(Fig 4C). Interestingly, as in the study by Schmack and colleagues (Schmack et al. 2013) this functional connectivity was related to the conviction scores for the delusion-prone group (Fig 5). Although this effect was also observed for the total PDI-scores, it remained when tested for the normalised convictions scores. Thus, the conviction scores had a specific effect on the connectivity between lOfc and right posterior insula independent on the number of endorsed delusional items.

**Fig. 5.**
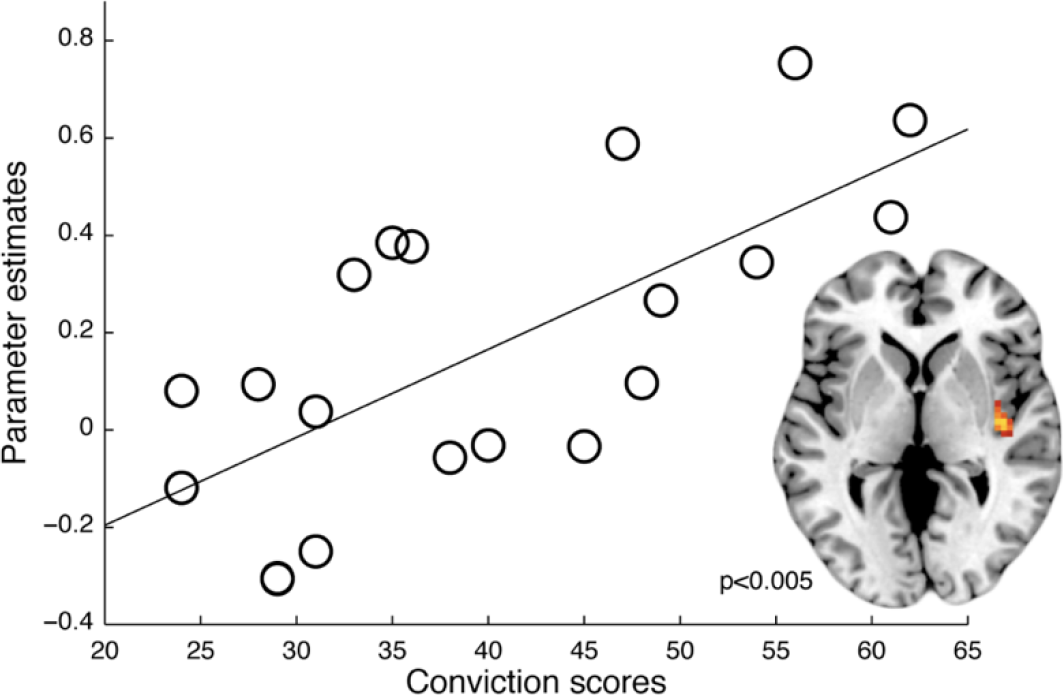
Relation between delusion-proneness and functional connectivity. The functional connectivity (PPI-analysis) between the right lOfc and i.e. right posterior insula ROI as an effect of instructions correlated with conviction scores in the delusion-prone group (Z = 3.44, pFWE = 0.003). A similar effect was shown for PDI-total scores (Z = 3.29, pFWE = 0.004) and normalised conviction scores (Z = 2.77, pFWE = 0.016).

The PPI-analysis of the effects of instruction on fear processing showed a significantly larger connectivity between the lOfc and the caudal anterior cingulate cortex (cACC), overlapping with fear related activation, in delusion-prone compared to control participants (Z=2.96, corrected p=0.012) (Fig 4D).

lOfc is tightly related to successful re-appraisal (Eippert et al. 2007; Golkar et al. 2012; Kanske et al. 2011; Wager et al. 2008) and the placebo effect in pain (Wager & Atlas 2015; Petrovic et al. 2010; Petrovic et al. 2002) and emotion (Petrovic et al. 2010; Petrovic et al. 2005). Both conditions involve a change in the underlying rules that relate to the interpretation of an emotional experience and the associated expectations. The significant group difference in lOfc functional connectivity - combined with no difference between the groups in the activation level related to fear processing - suggests mainly a difference in the re-appraisal effect between delusion-prone and control subjects.

A similar region in lOfc that links expectations to visual input (Bar 2003), was suggested to mediate belief congruent information to visual processing of the random dot kinetogram illusion related to delusion-proneness (Schmack et al. 2013). Based on these studies, we argue that lOfc may be important for construction of higher-order priors used more readily in delusion-proneness, especially in emotional and visual processes.

In a previous study on instructed fear conditioning (Atlas et al. 2016), an effect of instructions was observed in the dorsolateral prefrontal cortex (dlPFC), stretching towards ventrolateral PFC. Our main activation in the lOfc extends towards the same area. Finally, only the delusion-prone group showed activation in the ventromedial prefrontal cortex (vmPFC) in main effect of instructions - a region previously implicated in mediation of cognitive reappraisal (Wager et al. 2008).

From a predictive coding perspective the present study together with previous findings (Schmack et al. 2013; Teufel et al. 2015), suggest that individuals in psychosis-related states, including healthy delusion-prone subjects, are more prone to integrate and use higher-order beliefs (or models/priors) of the world in order to better comprehend a noisy perceptual environment. Our results are in sharp contrast to previous findings in studies on low-level bottom-up fear learning, such as classical fear conditioning, where only attenuated effects have been observed (Balog et al. 2013; Holt et al. 2009; Holt et al. 2012; Jensen et al. 2008; Romaniuk et al. 2010). Also, as we showed that these individuals integrate higher-order information more readily than controls even before the conditioning, simple adjustment to low-level aberrant salience (Kapur 2003) cannot solely explain overly stable beliefs. Altogether, our study and previous work on fear processing in psychosis-related states, suggest the coexistence of a weak low-level, and strong high-level fear learning in psychosis-related endophenotypes.

### Methods

#### Participants

We screened 925 male individuals aged 18 to 35 years (mean 24.98 years, SD 0.161) for delusion-proneness using *PDI (Peters’ Delusion Inventory* - 21 items) (Peters et al. 2004). The subjects also completed *ASRS (World Health Organization Adult ADHD Self-Report Scale)* (*2*), and *AQ (Autism Spectrum Quotient questionnaire)* (*3*) to control for sub-clinical tendencies of ADHD (Attention and Hyperactivity disorder) and ASD (Autism Spectrum disorder) (Louzolo et al. 2017). Participants were recruited through social media and filled in online versions of the questionnaires. It was stressed twice that they had to be healthy and without any psychiatric history. Upon submission of their contact details and after giving their consent, participants received a link to the questionnaires and an automatically generated unique ID-code that they used when filling in the questionnaires.

Based on the questionnaire results we selected 51 right-handed male individuals aged 18-35 years; out of which 26 were in the control group (PDI scores ranging from 2 to 6), and 25 in the delusion-prone group (PDI scores ranging from 10 to 17). Due to technical issues during the scanning procedures (movement and technical problems with the stimulation device), 8 participants had to be removed from both behavioural and imaging analyses. A total of 43 participants (control group: n=23, PDI mean=3.78, SD=1.38, and delusion-prone group: n=20, PDI mean=12.85, SD=1.84) thus underwent a successful *delayed fear conditioning* procedure in a 3T GE MR scanner and contributed to the behavioural results. Out of those 43 participants, another 3 were removed from the imaging analyses due to large movement artefacts, resulting in a total of 20 participants in each group contributing to the fMRI results (control group: PDI average =3.85 and SD =1.37; delusion-prone group: PDI average =12.85 and SD =1.84). The size of the two groups were comparable to previous fMRI studies on conditioning and psychosis related states (Balog et al. 2013; Holt et al. 2009; Holt et al. 2012; Jensen et al. 2008; Romaniuk et al. 2010).

All participants gave once again their informed consent before the experiment, and were paid 450 SEK for their participation. The study was approved by the regional ethical board of Stockholm (www.epn.se).

#### Stimuli and apparatus

In the conditioning paradigm the unconditioned stimulus (UCS) consisted of a mildly aversive electric stimulation. Prior to the start of the experiment a pair of Ag/AgCl electrodes (27 × 36 mm) was attached to participants’ left forearm with electrode gel and used to deliver electrical stimulation. Before lying down in the scanner, participants went through a standard work-up procedure, during which stimulation intensity was gradually increased until participants judged it as unpleasant, but not intolerably painful. Stimulus delivery was controlled by a monopolar DC-pulse electric stimulation (STM200; Biopac Systems Inc., www.biopac.com). Each electrical stimulation lasted for 200ms, co-terminating the presentation of the reinforced CS+ stimuli. The experiment was presented using Presentation (www.neurobs.com, version 9.13) and was displayed on a screen inside the scanner. Participants controlled the computer cursor through the use of a trackball device.

The paradigm started with an instruction phase that was followed by a fear acquisition phase, and ended with an extinction phase (Fig. 1A). The conditioned stimuli (CS) consisted of four Caucasian male faces (selected from a picture set used in Johansson *et al*) (4) displaying a neutral facial expression (2 CS+ and 2 CS-) and randomised between participants. For illustration purposes we used silhouettes on the timeline sketch Fig.1.

In the instruction phase two of the faces (instructed CS+ and CS-; iCS+/iCS-) were coupled with information about their contingencies with the UCS (including a fabricated short description about their personality and the risk of being associated with a “shock”). The two other CS faces (non-instructed CS+ and CS-; niCS+/niCS-) contained no information about their contingencies with the UCS. The phrasing used in the instructions is presented in Fig. 1B (original text in Swedish).

In the acquisition and extinction phases each CS was displayed 12 times for 5 seconds, and the jittered inter-trial interval was 11.5± 2 seconds. The CS+ were coupled with UCS with a 50% contingency in the acquisition phase and there was no UCS in the extinction phase.

#### Skin conductance response

Skin conductance was recorded during the whole session. Two Ag/AgCl electrodes (27 × 36 mm) were attached to the distal phalange of the first and third fingers of participants’ left hand. The skin conductance response (SCR) was amplified and recorded using an fMRI compatible BIOPAC Systems (Santa Barbara, CA). Data were analysed using AcqKnowledge software (BIOPAC Systems). Processing of the raw data consisted of low-pass (1Hz) and high-pass (0.05Hz) filtering. For each CS, the conditioned SCR amplitude was quantified as the peak-to-peak amplitude difference to the largest response, in the 0.5–4.5 sec latency window after the stimulus onset. The SCRs were transformed into microSiemens (μS), and responses below 0.02 μS were encoded as zero. A square-root transformation was applied to raw SCRs to normalise the data distribution. Participants who displayed a SCR to less than 20% of each of the two CS+ were considered non-responders and excluded from SCR analyses. This resulted in 18 controls and 20 delusion-prone participants that were used in the SCR analysis.

#### Behavioural analyses

Since our focus was on explicit learning we used evaluative fear measurements (Petrovic et al. 2008) as our main outcome. On several occasions throughout the experiment (before instructions, during instructions, before acquisition, before and after extinction) participants had to rate how friendly each CS looked, using a visual analogue scale with “the least sympathetic person you can imagine” stated on the left anchor, and “the most sympathetic person you can imagine” on the right anchor (originally in Swedish). The X-axis coordinates of the scale were converted into numbers, from −100 (left anchor) to +100 (right anchor) and used as the rating scores. The first rating of each CS was referred to as the baseline rating and used to normalise the subsequent ratings for a given CS. The normalised scores were computed for each CS, by subtracting the first ratings from the following ratings. In order to estimate learning in our paradigm we calculated the difference between CS- rating and CS+ rating, in each pair (instructed and non-instructed). This difference score is referred to as “*affective learning index*” and represents the main outcome value in the study as we were interested in explicit learning. Instructions were presented twice (followed by ratings: T1’ and T1) in order to increase explicit learning (Fig. 1A). Out of these two ratings we used the one following the second instruction presentation (T1) in subsequent analyses as it represented the total effect of the instruction manipulation. This resulted in four *affective learning indices:* 1) T0 - before instruction learning 2) T1 - after instruction learning, 3) T2 - after acquisition and 4) T3 - after extinction (Fig. 1A). During the debriefing session after the experiment, participants were also asked to rate how strongly they felt they had been influenced by instructions and aversive stimulation, respectively (0: no influence at all, 10: extremely high influence).

Two specific hypotheses were tested for the behavioural part of the study:

−*Main hypothesis:* As psychosis-proneness has been associated with stronger learning and use of high-level priors (Schmack et al. 2013; Teufel et al. 2015), instructions should have a greater influence on fear learning in the delusion-prone subjects than in the normal population. We therefore hypothesised that the delusion-prone group would show larger instructed *affective learning index* in all phases compared to the control group.

−*Secondary hypothesis:* In line with previous studies on fear conditioning (Romaniuk et al. 2010; Jensen et al. 2008; Holt et al. 2009; Holt et al. 2012) we hypothesised that delusion-prone individuals would display an attenuated conditioning effect. This would be reflected by significantly smaller non-instructed *affective learning index* following acquisition in the delusion-prone group as compared to controls. Another secondary hypothesis was that the extinction effect would be smaller in the delusion-prone group mirroring *bias against disconfirmatory evidence* described in psychosis-related states, whereby schizophrenia patients (Woodward et al. 2008; Moritz & Woodward 2006; Veckenstedt et al. 2011; Woodward et al. 2006; McLean et al. 2016) and delusion-prone individuals (Buchy et al. 2007; Woodward et al. 2007; Orenes et al. 2012) exhibit a tendency to disregard evidence that goes against the current assumption.

In summary, on a behavioural level we expected increased effect of instructions on fear learning (instructed fear learning) but decreased effects of normal fear conditioning and a lesser extinction effect associated with delusion-proneness. We used one tailed t-tests for hypothesized and predicted results (indicated in text), and two-tailed t-tests otherwise.

#### Functional Imaging analysis

We hypothesized that lateral orbitofrontal cortex would have a decisive role in the increase of fear learning due to instructions - based on its previously shown involvement in processes where expectations have been experimentally manipulated. Studies on cognitive reappraisal have suggested that lateral orbitofrontal cortex (lOfc) is specifically involved in reappraisal success (Eippert et al. 2007; Golkar et al. 2012; Kanske et al. 2011; Wager et al. 2008). Moreover, it has been suggested that the lOfc is specifically involved in the expectation effect of placebo analgesia (Wager & Atlas 2015; Petrovic et al. 2010; Petrovic et al. 2002) and emotional placebo (Petrovic et al. 2010; Petrovic et al. 2005). Data from these studies suggests that the right lOfc, especially, is involved in placebo (Petrovic et al. 2010; Petrovic et al. 2002; Petrovic et al. 2005) and cognitive reappraisal processes (Wager et al. 2008).

We argue that instructed fear learning is linked to both higher order expectation effects and cognitive reappraisal in pain and emotion, and we therefore examined the acquisition phase results with a primary focus on effects in lOfc. Further, we posited that any behavioural effects in relation to instructed fear learning in the delusion-prone group would be linked to functional or effective connectivity effects in the right lOfc as previously observed in cognitive reappraisal (Wager et al. 2008).

Apart from the general hypothesis about the involvement of lOfc in the instruction effects, we more specifically hypothesized that the delusion-prone group would exhibit (i) increased lOfc responses to instructed fear learning, and (ii) increased effective connectivity between the lOfc, and pain and fear regions as an underlying mechanism associated with a stronger effect of instructions on *affective learning index*.

Due to limited space, we constrained the present functional imaging analysis to the acquisition phase.

#### Image acquisition

Participants were scanned in a 3T MR General Electric scanner with a 32-channel head coil. A T1-weighted structural image was acquired before the beginning of the paradigm. Functional scans were obtained using a gradient echo sequence T2*-weighted echo-planar imaging (EPI) scan (TR=2.334 sec, TE=30 ms, flip angle=90 degrees, 49 axial slices in ascending order (thickness=3 mm) and a field of view (FOV)=22cm, matrix size=72×72−3mm). The first four scans were defined as dummy scans and discarded from the analysis. Functional image acquisition comprised 2 runs of 245 volumes each (acquisition and extinction phases, respectively), with a break of approximately 4-5 minutes between them.

#### Imaging data analysis

Data pre-processing and analyses were performed using a default strategy in the SPM8 software package (Statistical parametric mapping, Welcome Department of Cognitive Neurology, London, UK http://www.fil.ion.ucl.ac.uk/spm). For each participant, individual images were first slice-time corrected and realigned to the first volume to correct for head movement. The T1-weighted image was then co-registered with the mean EPI image, segmented and normalised to the Montréal Neurological Institute standard brain (MNI). Then, functional images were spatially smoothed with an 8-mm full width at half maximum (FWHM) isotropic Gaussian kernel, and a temporal high-pass filter with a cut-off of 128 seconds was used to remove low-frequency drifts.

A general linear model (GLM) comprising 9 regressors was defined at the first-level analysis; one regressor per CS type (iCS+, iCS-, niCS+ and niCS-) with each onset modelled as a 5-second event, and one regressor for the UCS presentation. In addition, these four regressors (excluding UCS) were also parametrically modulated with a linearly changing function to capture activity changes over time. All 9 regressors were convolved with the canonical hemodynamic response function and entered into the GLM as implemented in SPM. Motion regressors were also included in the model. The two phases of the experiment (acquisition and extinction) were modelled and analysed separately.

We first analysed main effects of fear (CS+ vs. CS-) and the interactions with instructions. Similarly, we examined the main effects of pain. We also analysed possible differences between delusion-prone and control groups in these activations using a ROI approach in order to increase the sensitivity. A small volume correction in a spherical ROI (6 mm radius) was then applied in the contrasts between the two groups. The ROIs were centred over the maximally activated voxels in caudal ACC (cACC) and anterior insula in the main effect of fear and in posterior insula in the main effect of pain. The results were assessed at p<0.05, family-wise error (FWE) corrected for multiple comparisons.

To test our main hypotheses regarding the functional imaging results, we first conducted a GLM group analysis to compare the effect of instruction in the lOfc for delusion-prone compared to control participants. The results were assessed at p<0.05, family-wise error (FWE) corrected for multiple comparisons. Given our a priori hypothesis, we used small-volume correction (SVC) for multiple comparisons within an anatomical lOfc ROI defined using the pick atlas in the SPM, in addition to whole brain analysis.

We also examined effective connectivity using a psychophysiological interaction (PPI) analysis in SPM (Friston et al. 1997). This analysis identifies context-induced changes in the strength of connectivity between brain regions, as measured by a change in the magnitude of the linear regression slope that relates their underlying neuronal responses. Significant PPI results indicate that the contribution of one area to another changes with the experimental context (Friston et al. 1997). We assessed connectivity changes between the right lOfc and the rest of the brain. The lOfc seed region was defined using a sphere with a radius of 6 mm centered on the right lOfc group maximum from the GLM analyses of instruction-related activity. For each participant, the seed was adjusted to center on the individual peak response within the group seed sphere, and the fMRI time series was extracted and deconvolved to generate the neuronal signal. We then conducted two PPI analyses using the contrast (i) instructed vs. non-instructed [(iCS+ and iCS-) vs (niCS+ and niCS-)] and (ii) the interaction effect (fear learning in instructed vs. fear learning in non-instructed stimuli; [(iCS+ vs iCS-) vs (niCS+ vs niCS-)]) as the psychological factor. For each participant, a GLM was conducted including three regressors representing the time course of the seed region (the physiological factor), the psychological factor and their product (the PPI). The parameter estimates for the PPI regressor from each participant were then entered into a second-level analysis, and we again assessed the results at pFWE<0.05.

We conducted SVC in several ROIs for the PPI analyses. First, we used the group-level main effect of fear learning (CS+ vs. CS-) to identify cACC and anterior insula (Table S1). Second, we examined any group differences in low-level sensory processing areas, in line with previous findings of altered effective connectivity between the lOfc and the visual cortex (Schmack et al. 2013). To obtain a low-level sensory region, we used the group-level main effect pain (mildly painful electric stimulation) to identify the posterior insular cortex. This region has been the most consistently reported brain activation site across all pain conditions and is considered a nociceptive input area (Tanasescu et al. 2016).

Finally, we assessed whether there was a significant correlation between conviction scores and the functional connectivity between the lOfc seed region and low-level sensory regions (i.e. defined as posterior insular in the present study) to investigate whether we could reproduce the findings by Schmack and colleagues (Schmack et al. 2013). On a more exploratory level, we analysed whether such a correlation was also present for the total PDI-score, the normalised conviction score as well as the two other sub-scores in PDI (distress score and preoccupation scores).

#### Source data

Behavioral source data is provided for Figure 2-3: S Figure 1-5 and can be found on: http://doi.org/10.5281/zenodo.1170599.

## Supplementary Materials

**Fig. S1.**
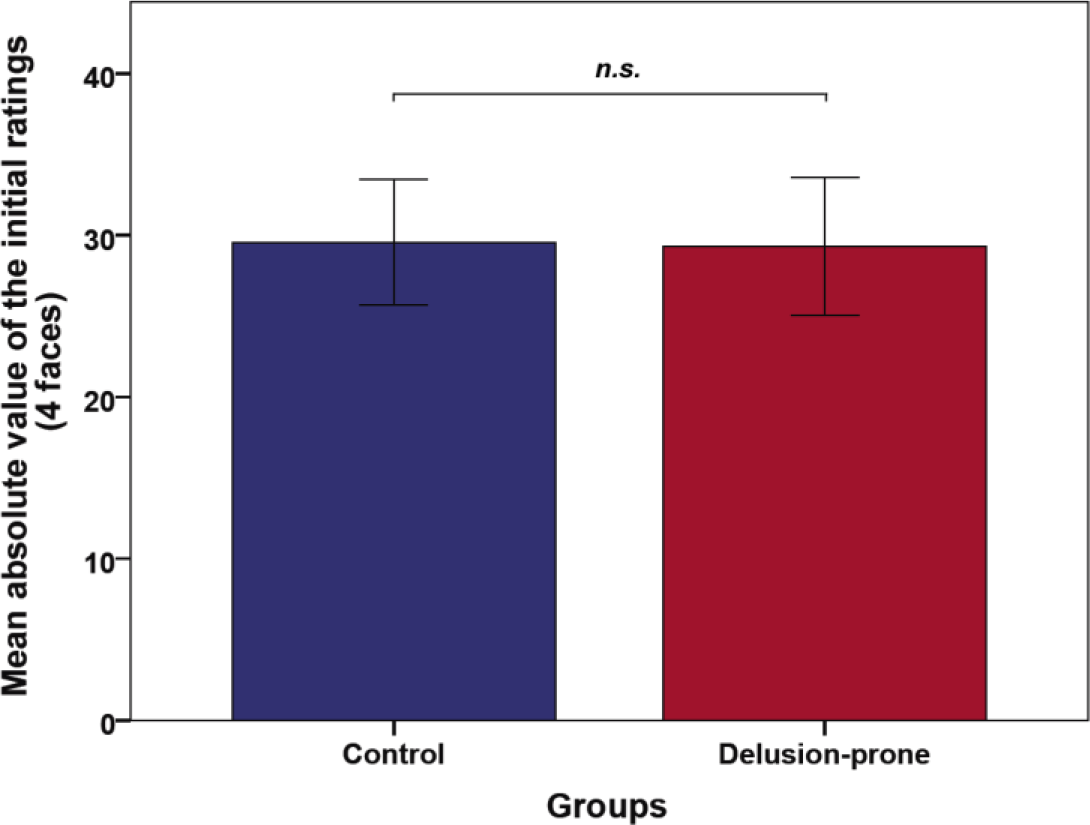
Baseline ratings of the four faces used in the experiment. A baseline rating for each face was performed before any information was presented and used for normalisation of subsequent ratings. We first ran a one-way ANOVA to confirm there was no significant difference between the ratings of the four faces. The ANOVA was not significant (F(1,170)=1.420, p=0.239), suggesting that the four faces did not differ in terms of initial ratings. We then tested whether groups (control group and delusion-prone group) differed on the averaged absolute value of the initial ratings, and found no significant difference (t=0.082, p=0.936, independent two-sample t-test). This result suggests that group differences associated to instructions or conditioning cannot be explained simply by a difference between the groups in their general rating strategy. **Error bars: 2 S.E.**

**Fig. S2.**
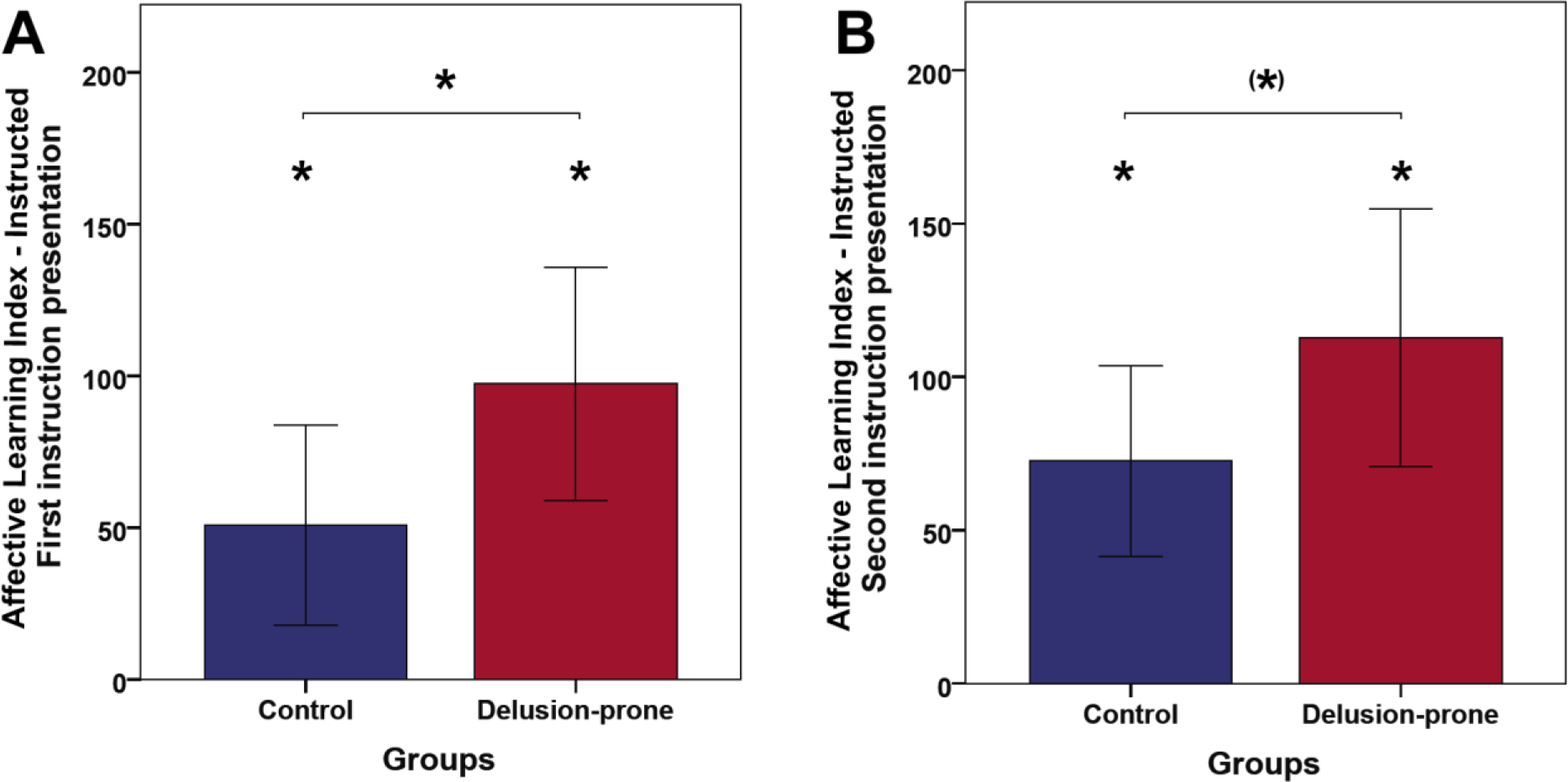
Effect of the (A) first and (B) second instruction presentations on *affective learning index*. The *affective learning index* showed a clear learning effect after the first instruction phases for the instructed stimuli (iCS+/iCS-) [*control group*: mean=50.65, SD=79.06, one-sample t test t=3.073, df=22, p=0.003 one-tailed; *delusion-prone group:* mean=97.40, SD=85.89, one-sample t test t=5.07, df=19, p<0.001 one-tailed]. The group difference was significance (independent t-test t=-1.858, df=41 p=0.035 one-tailed). In both groups the *affective learning index* increased significantly after the second instruction presentation (*control group:* mean=72.52, SD=74.59, paired t-test t=-1.963; df=22, p=0.032 one-tailed; *delusion-prone* mean=114.75, SD=93.26, paired t-test t=-2.350, df=19, p=0.015 one-tailed). The group difference was on the border of significance (independent t-test t=− 1.649, p=0.053 df=41 one-tailed). **Error bars: 2 S.E.**

**Fig. S3.**
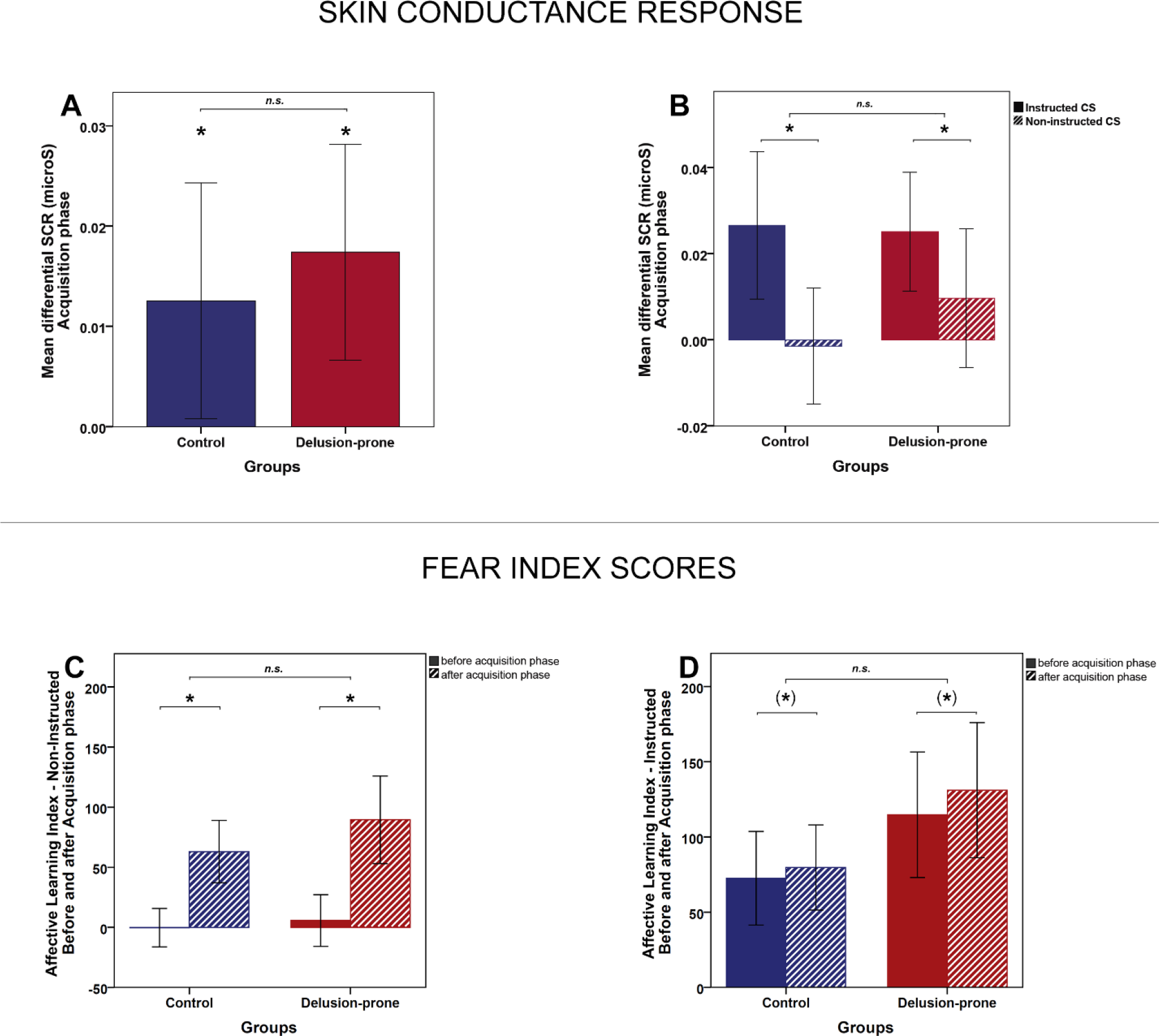
Effect of fear conditioning on skin conductance and *affective learning index* in each group. The t-test on the differential SCR (SCR-CS+ vs SCR-CS-) was significantly different from zero for the acquisition phase (average=0.0151, SD=0.0271; t=3.424, df=37, p=0.001 one-tailed) suggesting a significant conditioning effect for all subjects. **(A)** This was also the case for each group (controls mean=0.0126μS, SD=0.0248, one-sample t-test t=2.145, df=17, p=0.024 one-tailed-delusion-prone mean=0.0174μS, SD=0.0296, one-sample t-test t=2.628, df=19, p=0.009 one-tailed). There was no group difference (independent two-sample t-test t=-0.741, df=73, p=0.461). During the extinction phase the differential SCR was no longer significantly different from zero (one-sample t-test t=-1.115, df=75, p=0.268). **(B)** In both groups, the differential SCR was mainly driven by the instructed CS-pair as suggested by a significant difference between the instructed and non-instructed condition (controls instructed mean= 0.0266μS, SD=0.036, non-instructed mean=-0.015μS, SD=0.029; paired t-test t=2.780, df=17, p=0.014 - delusion-prone instructed mean= 0.0251μS, SD=0.031, non-instructed mean=0.010μS, SD=0.036; paired t-test t=2.188, df=19, p=0.042). However, there was no significant interaction between the groups. Overall, it should be noted that the SCR data recorded in the fMRI scanner was noisy. We only used participants who showed a SCR to at least 20% of the presentations of each CS (hence, considered as responders), however many of them were characterised by a low reactivity. **(C)** *Affective learning index* for the non-instructed pair increased significantly after acquisition (T2 vs T1), however there was no group interaction (control mean before acquisition=-0.261, SD=38.46; mean after acquisition=63.00, SD=62.16; paired t-test t=-4.405, df=22, p=0.000 one-tailed – delusion-prone mean before acquisition=5.70, SD=47.92; mean after acquisition=89.45, SD=81.52; paired t-test t=-6.165, df=19, p=0.000 one-tailed). **(D)** A trend towards a conditioning effect was also observed in both groups for the instructed *affective learning index* (control mean before acquisition=72.52, SD=74.59; mean after acquisition=79.73, SD=67.93; paired t-test t=-1.679, df=22, p=0.054 one-tailed – delusion-prone mean before acquisition=114.75, SD=93.26; mean after acquisition=131.15, SD=100.35 – paired t-test t=-1.704, df=19, p=0.053 one-tailed). There was also no group interaction. Finally, the total *affective learning index* after conditioning (T2) was larger for the instructed than the non-instructed conditions in the delusion-prone group (mean instructed *affective learning* index=131.15, SD=100.35; mean non-instructed *affective learning* index=89.45, SD=81.52; paired t-test t=2.198, df=19, p=0.041). However, this was not the case in the control group (mean instructed *affective learning* index=79.74, SD=67.93; mean non-instructed *affective learning* index=63.00, SD=62.16; paired t-test t=1.000, df=22, p=0.328), although there was no significant difference of these effect between groups. Error bars: 2 S.E.

**Fig. S4.**
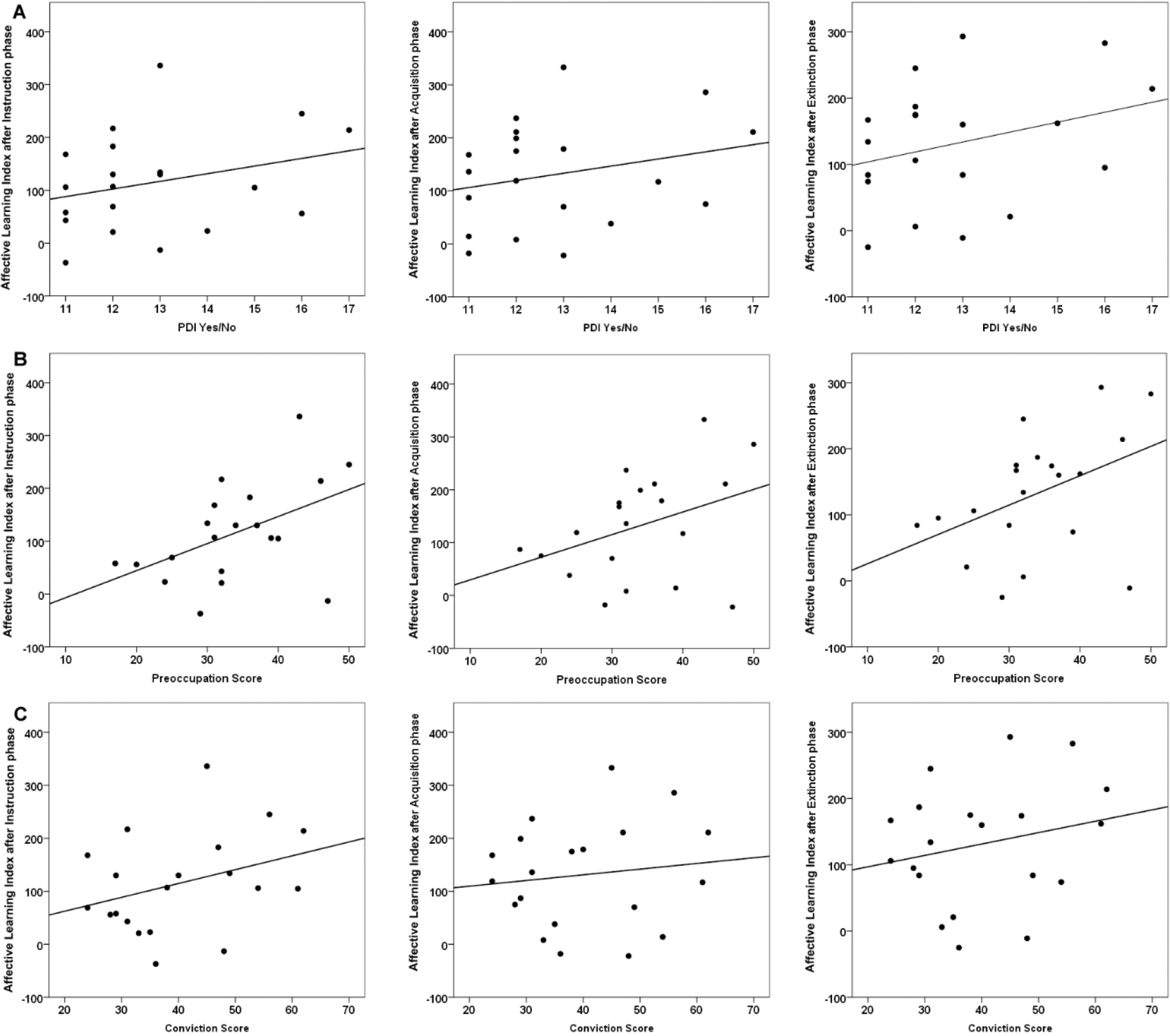
Correlations between instructed *affective learning index* and total PDI score, preoccupation score, and conviction score in delusion-prone individuals. No significant correlations were observed between *affective learning index* and the **total PDI score** (T1 (after instructions): r=0.255, p=0.277; T2 (after acquisition): r=0.224, p=0.342; T3 (after extinction) r=0.277, p=0.236, the **preoccupation score** (T1 (after instructions): r=0.442, p=0.051; T2 (after acquisition): r=0.386, p=0.093; T3 (after extinction): r=0.435, p=0.055, Pearson correlation tests), the **conviction score** (T1 (after instructions): r=0.282, p=0.230; T2 (after acquisition): r=0.081, p=0.733; T3 (after extinction): r=0.131, p=0.581, Pearson correlation tests) in the delusion-prone group. No correlation effects were observed for the control group. The equivalent correlation effects for distress effects are shown in main article.

**Fig. S5.**
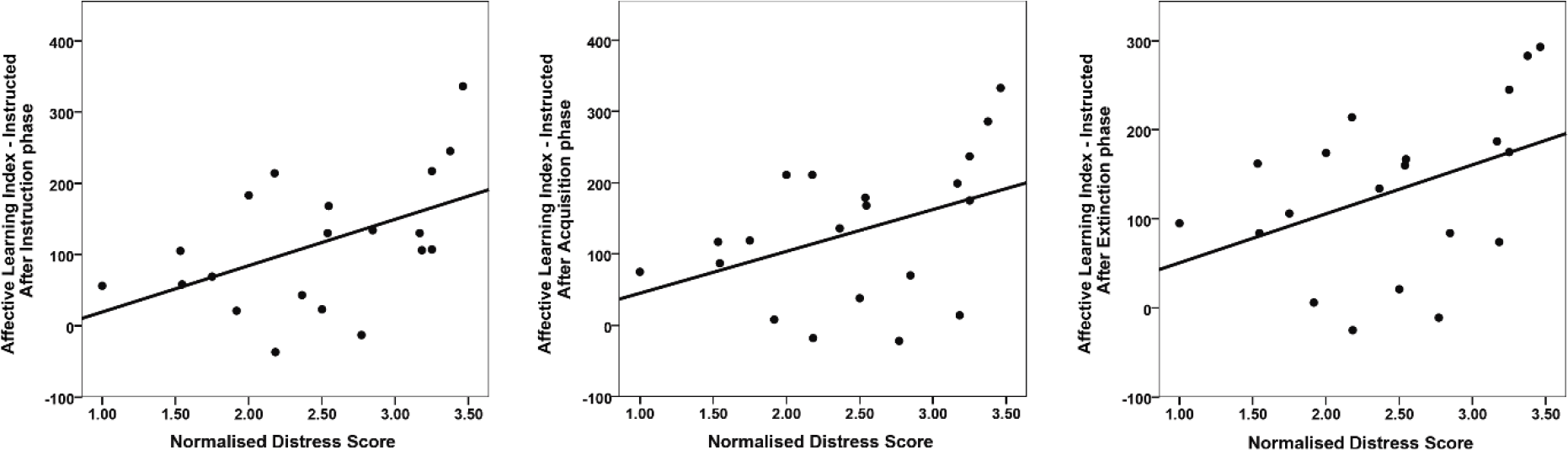
Relation between normalised distress scores and instructed learning. Since distress seemed as an important variable in relation to effects of instructions in our fear learning paradigm we explored it further. Only analysing the total sum of each of these subscores without taking the Yes/No score can be somewhat misleading, as it makes it difficult to differentiate between people who would score high on distress because they have a few delusion-like experiences that are extremely distressing, from people who score as high on distress because they have many delusion-like experiences that are not distressing at all. Normalising for the number of Yes/No scores provides a better estimate of how distressed, preoccupied and convinced participants are, unrelated to whether there is one or several delusion-like experience. We therefore compared the control and delusion-prone group in terms of sub-scores and found that the delusion-prone group had an average normalised distress score that was significantly higher than the control group (control=1.95, delusion-prone=2.47, t=-2.593, p=0.013, df=41, independent sample t-tests). There was no significant group difference in terms of normalised preoccupation or conviction scores. The normalised **distress scores** also correlated positively with *affective learning index* after the instruction phases in the delusion-prone group (r=0.527, p=0.017, Pearson correlation tests). This correlation reached a trend level in the acquisition and extinction phases (r=0.400, p=0.080; r=0.438, p=0.053, respectively - Pearson correlation tests, two-tailed). This normalised distress score was also positively correlated with the explicit rating of the influence of instructions, provided by the subjects after the experiment (r=0.491, p=0.028 Pearson correlation tests, two-tailed). No significant correlations were found in the control group.

**Fig. S6.**
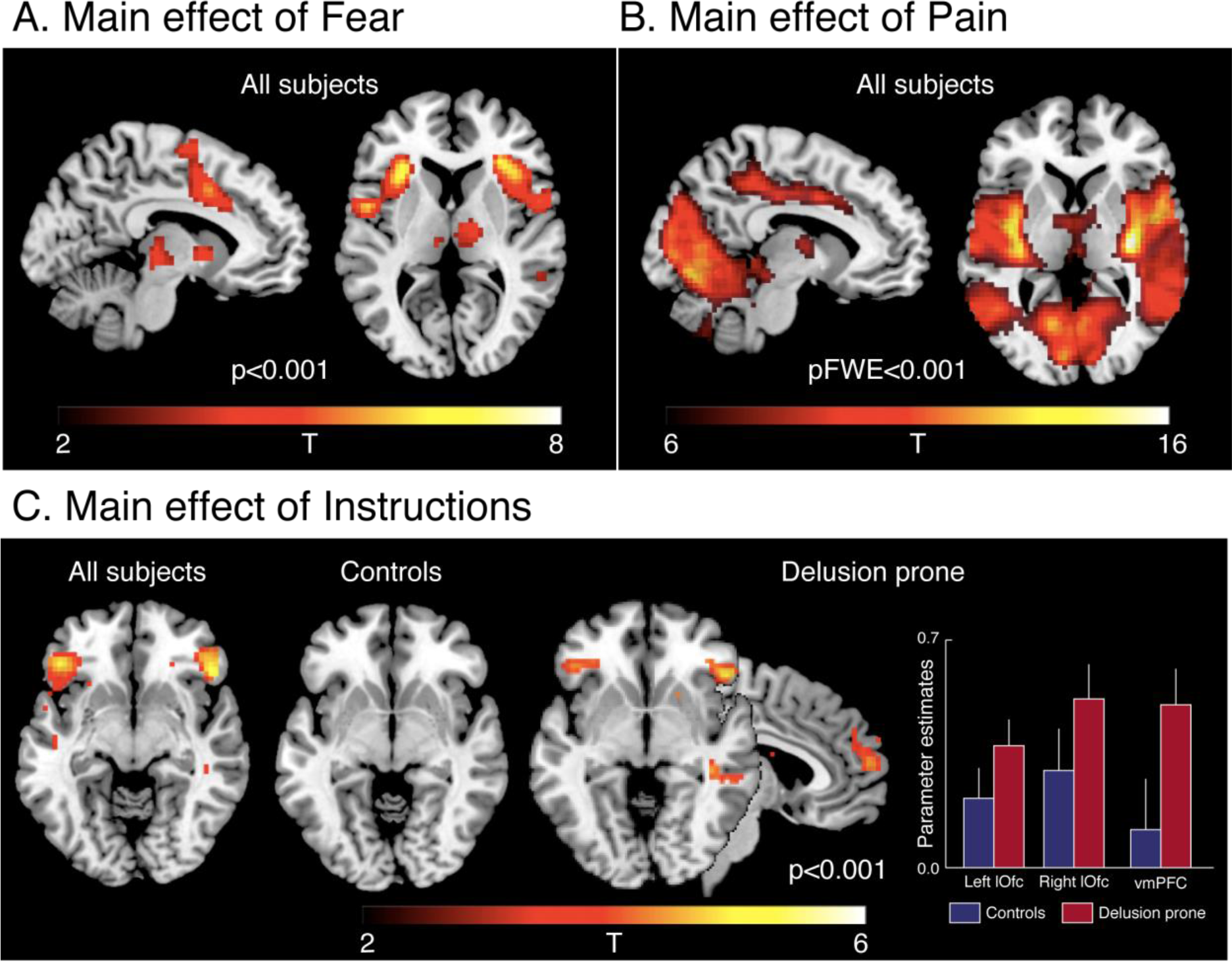
GLM results - whole-brain analyses. **(A)** The main effect of fear *(CS+ vs. CS-)* in all subjects led to activations in brain areas that are consistently reported in fear conditioning studies, including bilateral anterior insula, caudal anterior cingulate cortex (cACC), premotor/dorsolateral prefrontal cortex (dlPFC), right temporo-parietal junction (rTPJ) (Table S1). **(B)** The main effect of pain (shocks) in all subjects led of activation in brain areas associated with pain processing: caudal ACC (cACC), bilateral mid- and posterior insula - with the most pronounced activation in the left posterior insula (Table S5). However, as there was no control condition for pain, the activation pattern observed in this contrast was much wider than the one generally observed in pain studies using a control condition for pain. **(C)** The main effect of instructions showed a bilateral activation in lateral orbitofrontal cortex (lOfc) that was mainly driven by the delusion-prone group (there was no tendency of lOfc activation at the present threshold for the control group, while the lOfc activation was highly significant and bilateral for the delusion-prone group). In addition, delusion-prone individuals also displayed activation in the vmPFC that was not reported in the control group, nor in the all-subject activations (Table S2). Error bars: S.E.

**Fig. S7.**
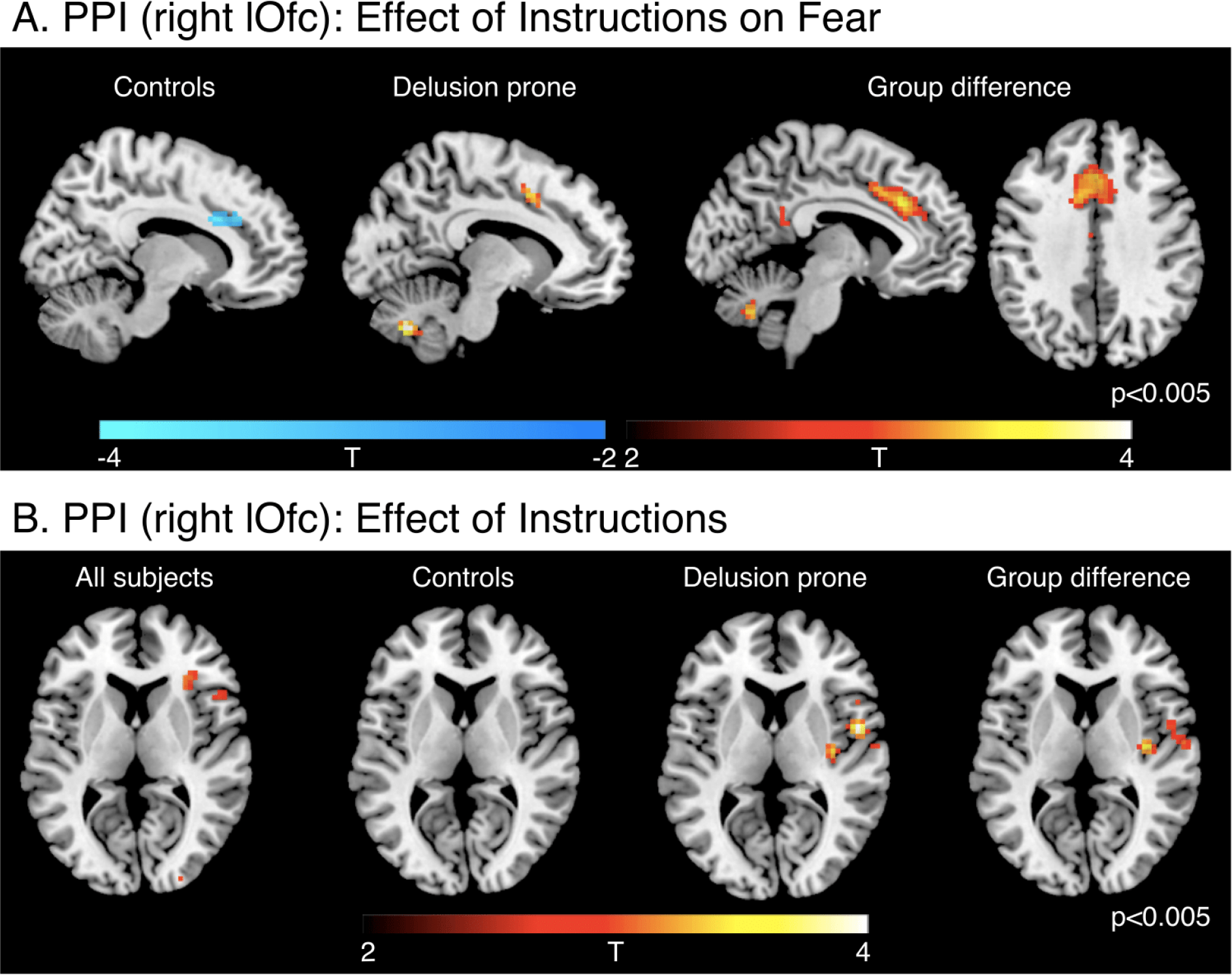
PPI results. **(A)** PPI-analysis of the effects of instruction on fear processing showed a significantly larger connectivity between lOfc and cACC, in delusion-prone compared to control participants (Z = 2.96, corrected p = 0.012). This effect was driven by a decreased connectivity in the control participants and an increased connectivity in the delusion-prone participants. **(B)** The PPI analysis also revealed an increased connectivity between the right lOfc and the anterior insular cortex in main effect of instructions, and a group difference in connectivity with the right posterior insula (Z = 3.29, corrected p = 0.004). This group difference was driven by the presence of a positive connectivity in the delusion-prone group. Error bars: S.E.

**Table S1.**
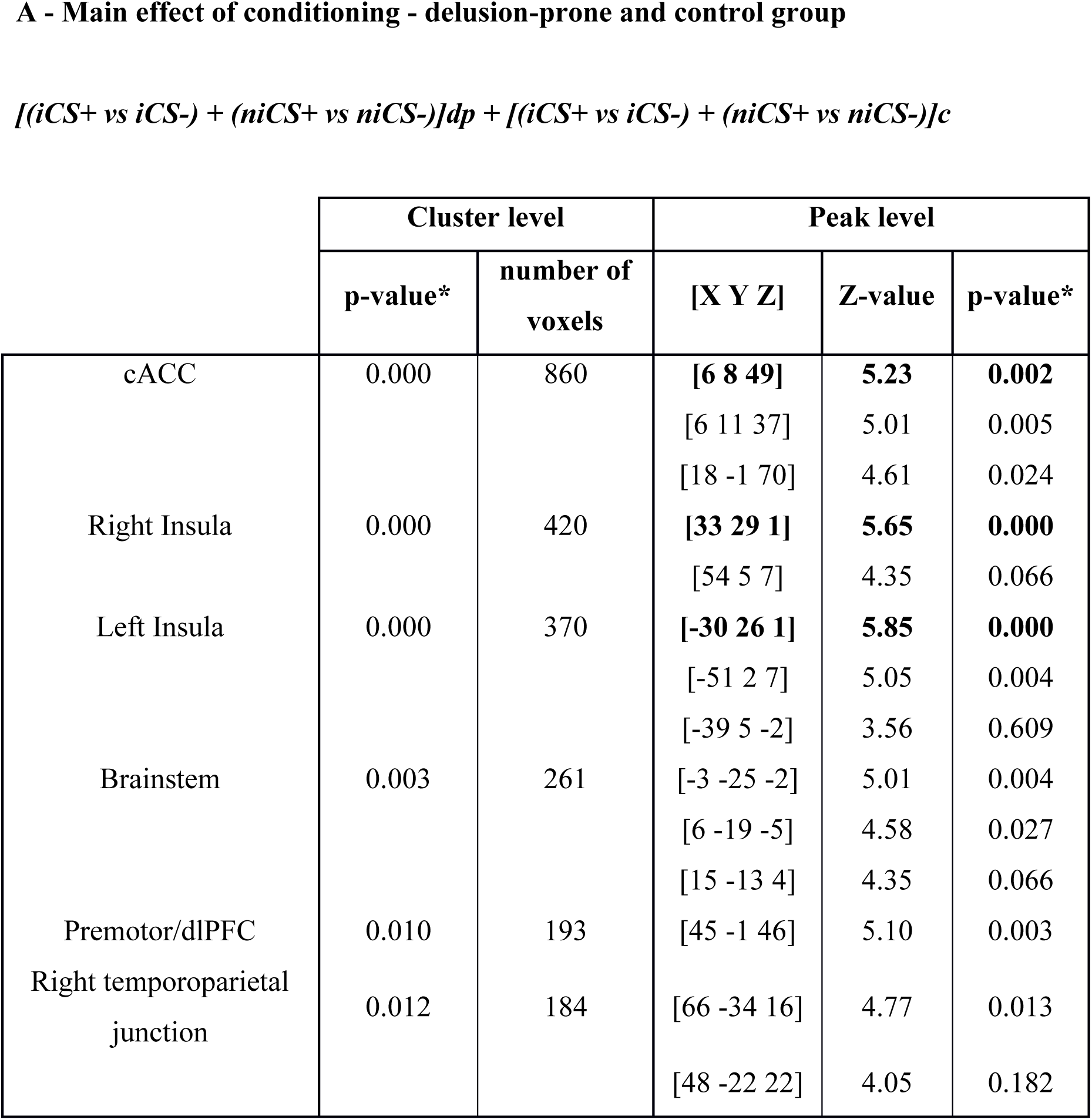
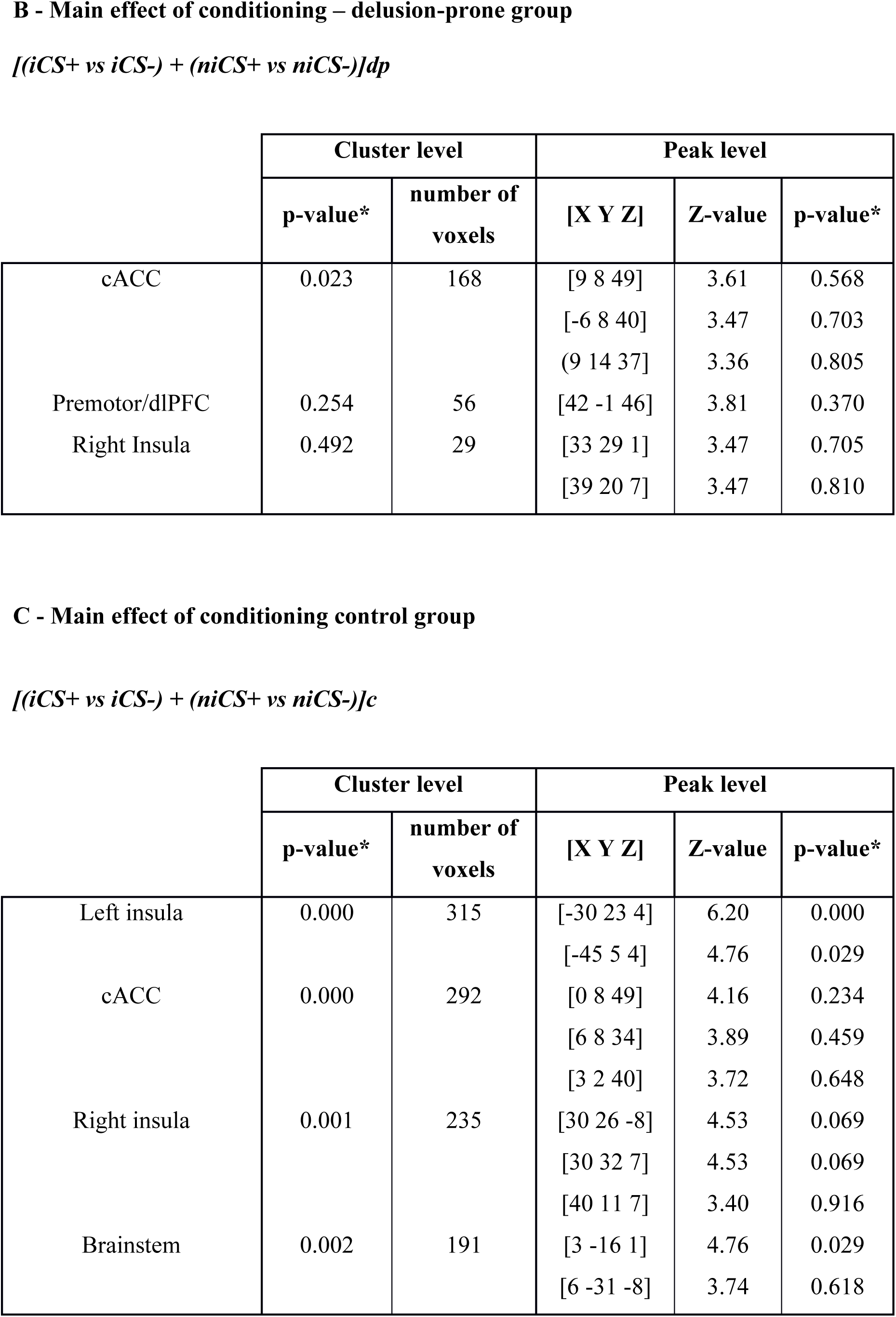
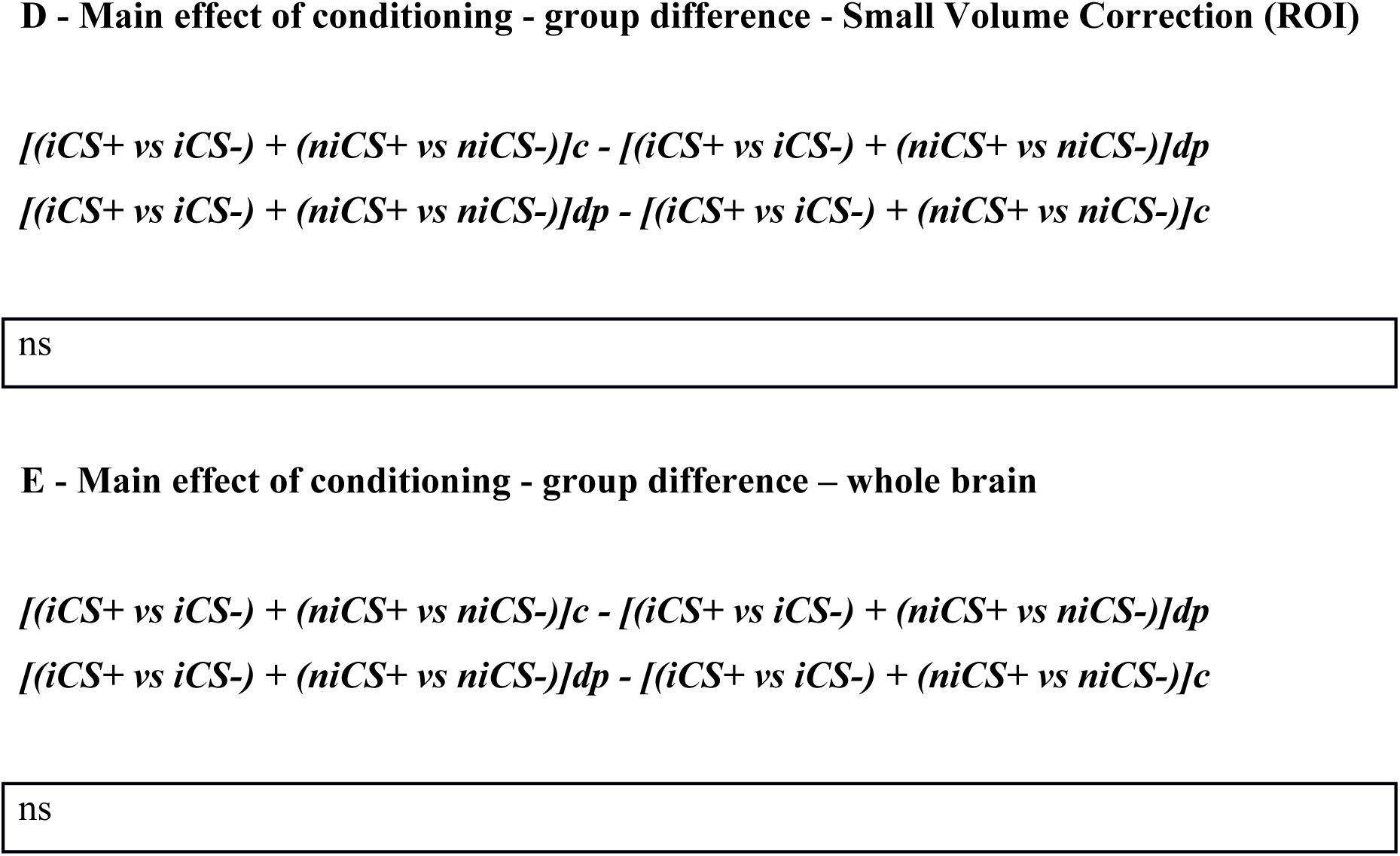
Effect of Conditioning. The main effect of the conditioning task *(CS+ > CS-)* led to activations in brain areas that are consistently reported in fear conditioning studies (22). The map was thresholded at p < 0.001 (uncorrected), k > 20 transformed voxels. dp: delusion-prone group. c: control group. The shown *p-values were corrected for full-brain volume (FWE-correction). Peak-activations used for the subsequent ROI-analyses (6mm sphere - including PPI-analysis) are shown in bold. cACC = caudal anterior cingulate cortex. dlPFC = dorsolateral prefrontal cortex.

|  | Cluster level |  | Peak level |  |  |
| --- | --- | --- | --- | --- | --- |
|  | p-value* | number of voxels | [X Y Z] | Z-value | p-value* |
| cACC | 0.000 | 860 | **[6 8 49]** | **5.23** | **0.002** |
|  |  |  | [6 11 37] | 5.01 | 0.005 |
|  |  |  | [18 -1 70] | 4.61 | 0.024 |
| Right Insula | 0.000 | 420 | **[33 29 1]** | **5.65** | **0.000** |
|  |  |  | [54 5 7] | 4.35 | 0.066 |
| Left Insula | 0.000 | 370 | **[-30 26 1]** | **5.85** | **0.000** |
|  |  |  | [-51 2 7] | 5.05 | 0.004 |
|  |  |  | [-39 5 -2] | 3.56 | 0.609 |
| Brainstem | 0.003 | 261 | [-3 -25 -2] | 5.01 | 0.004 |
|  |  |  | [6 -19 -5] | 4.58 | 0.027 |
|  |  |  | [15 -13 4] | 4.35 | 0.066 |
| Premotor/dlPFC | 0.010 | 193 | [45 -1 46] | 5.10 | 0.003 |
| Right temporoparietal junction | 0.012 | 184 | [66 -34 16] | 4.77 | 0.013 |
|  |  |  | [48 -22 22] | 4.05 | 0.182 |
|  | p-value* | number of voxels | [X Y Z] | Z-value | p-value* |
| cACC | 0.023 | 168 | [9 8 49]<br>[-6 8 40]<br>(9 14 37) | 3.61<br>3.47<br>3.36 | 0.568<br>0.703<br>0.805 |
| Premotor/dlPFC | 0.254 | 56 | [42 -1 46] | 3.81 | 0.370 |
| Right Insula | 0.492 | 29 | [33 29 1]<br>[39 20 7] | 3.47<br>3.47 | 0.705<br>0.810 |

|  | Cluster level |  | Peak level |  |  |
| --- | --- | --- | --- | --- | --- |
|  | p-value* | number of voxels | [X Y Z] | Z-value | p-value* |
| Left insula | 0.000 | 315 | [-30 23 4]<br>[-45 5 4] | 6.20<br>4.76 | 0.000<br>0.029 |
| cACC | 0.000 | 292 | [0 8 49]<br>[6 8 34]<br>[3 2 40] | 4.16<br>3.89<br>3.72 | 0.234<br>0.459<br>0.648 |
| Right insula | 0.001 | 235 | [30 26 -8]<br>[30 32 7]<br>[40 11 7] | 4.53<br>4.53<br>3.40 | 0.069<br>0.069<br>0.916 |
| Brainstem | 0.002 | 191 | [3 -16 1]<br>[6 -31 -8] | 4.76<br>3.74 | 0.029<br>0.618 |

|  |
| --- |
| ns |

|  |
| --- |
| ns |

**Table S2.**
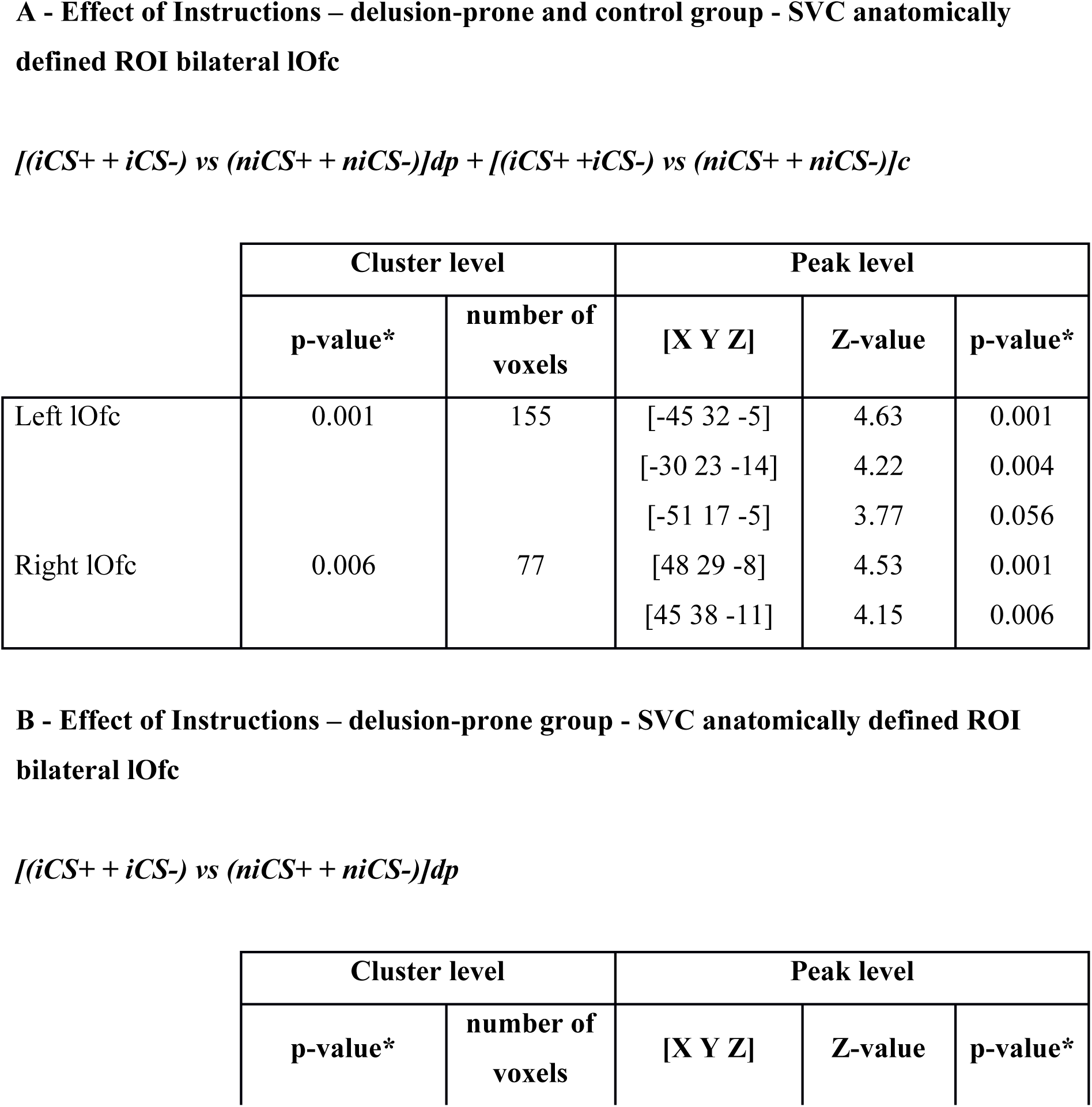
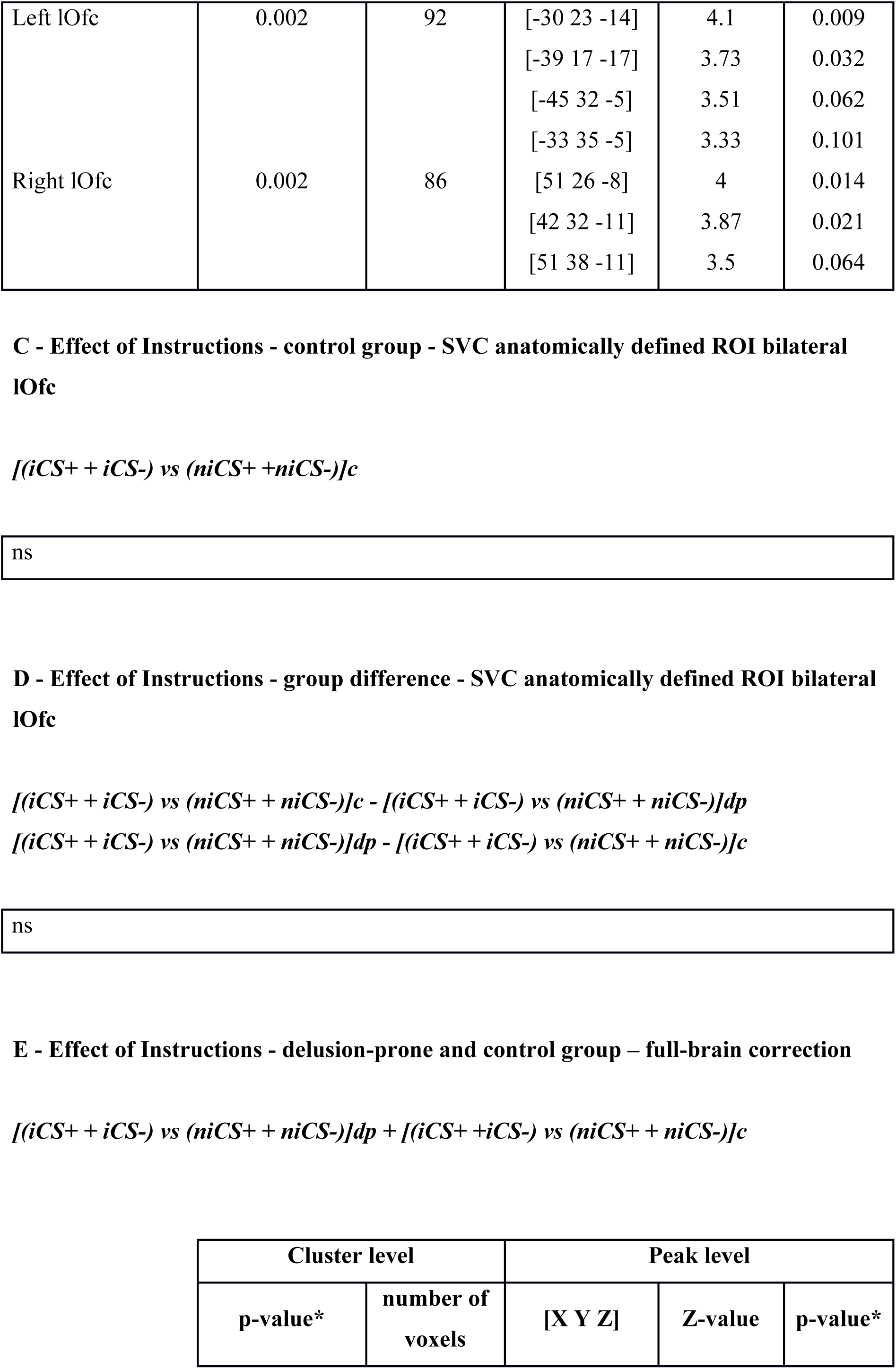
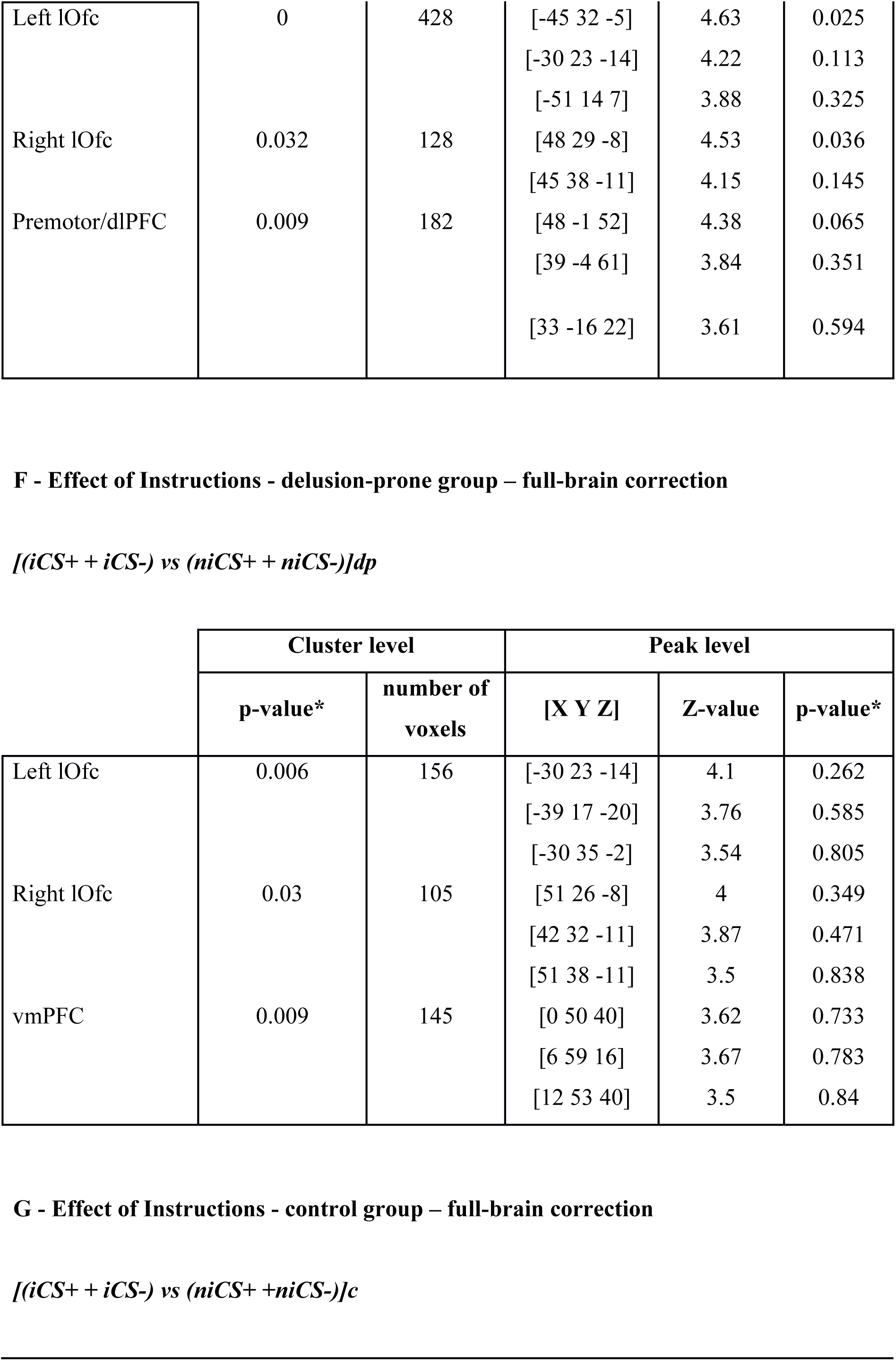
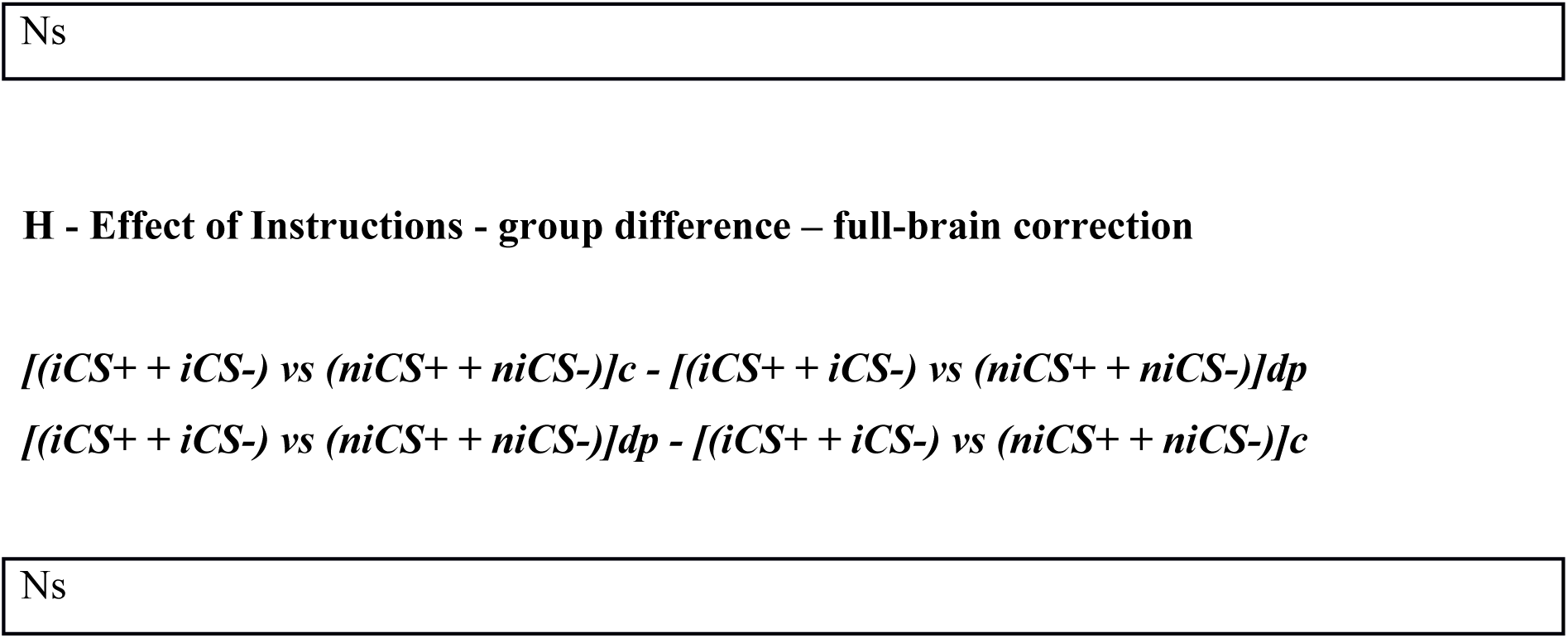
Effect of Instructions ROI analysis (A-D) and whole-brain analysis (E-H). The main effect of instructions resulted in the activation of brain regions involved in reappraisal conditions, i.e. lOfc bilaterally. The small volume correction analyses (A-D) were performed using an anatomically defined bilateral lOfc ROI. Importantly, the effects of learning did not yield any significant or non-significant activation in the control group (C), while the effects survived full-brain correction in the delusion-prone group (F). The general activation pattern (A and E) seems to be mainly driven by the delusion-prone individuals. The map was thresholded at p < 0.001 (uncorrected), k > 20 transformed voxels. dp: delusion-prone group. c: control group. The shown *p-values were corrected for full brain volume (FWE-correction). SVC = small volume correction.

**Table S3.**
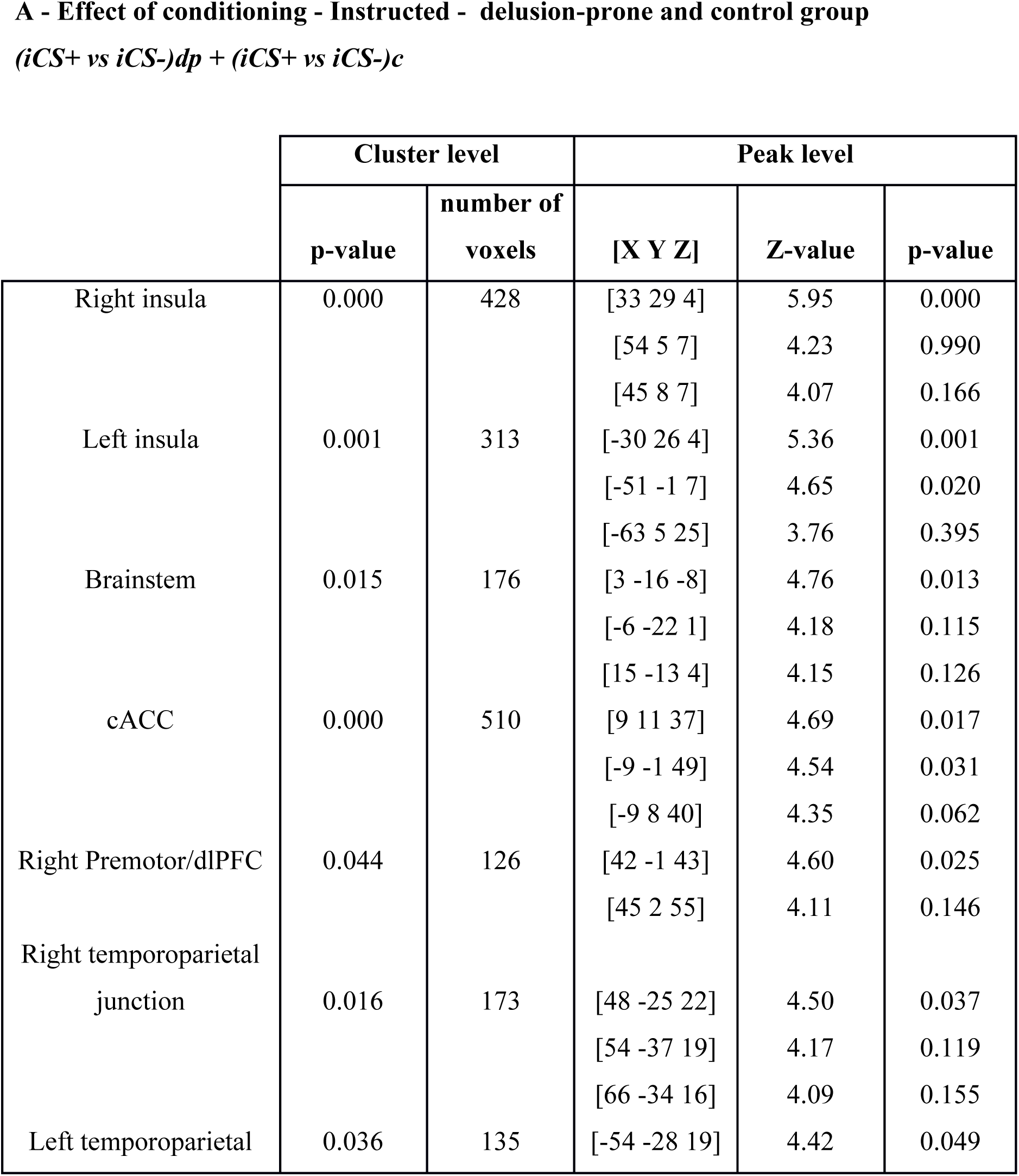
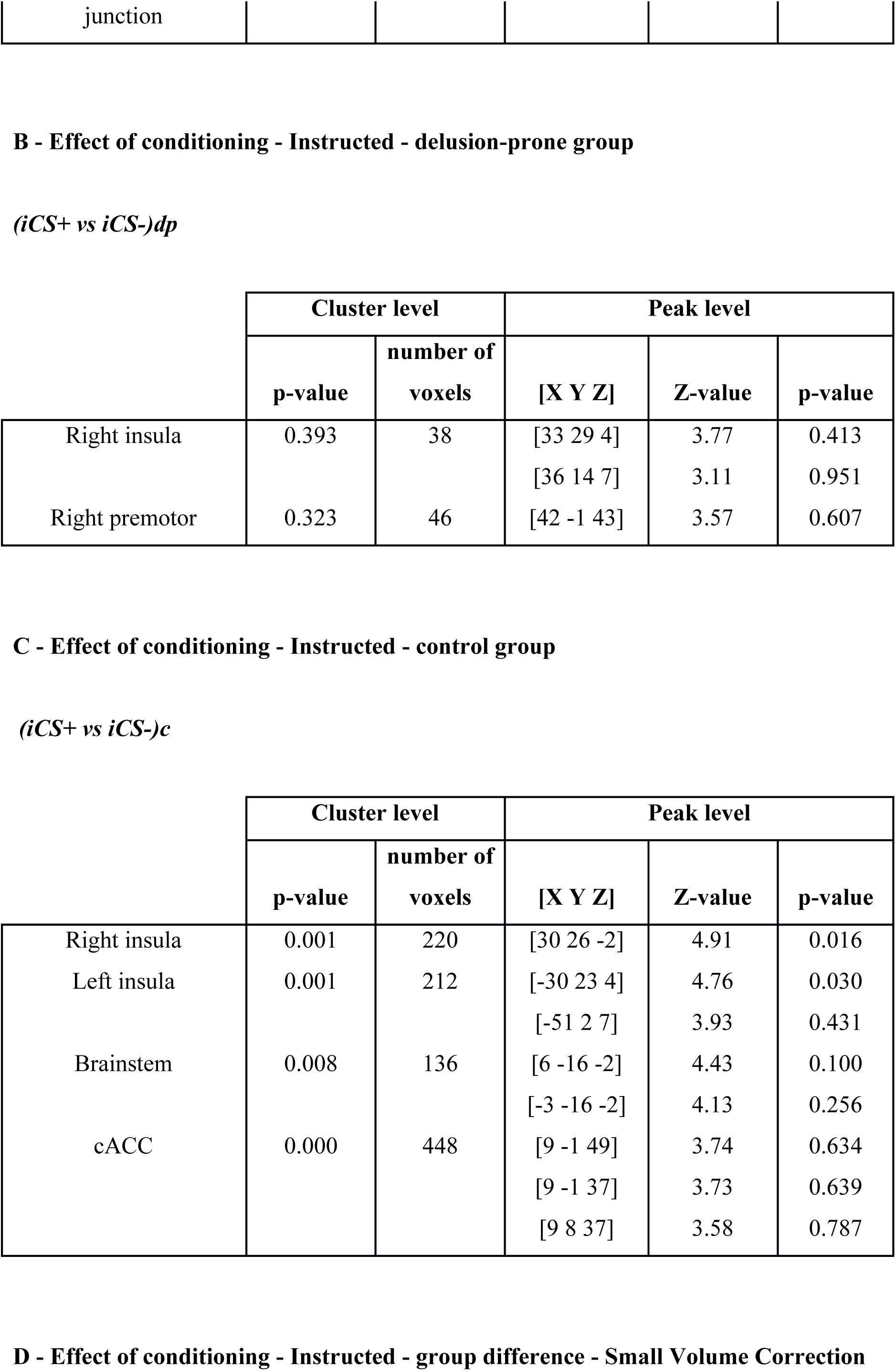
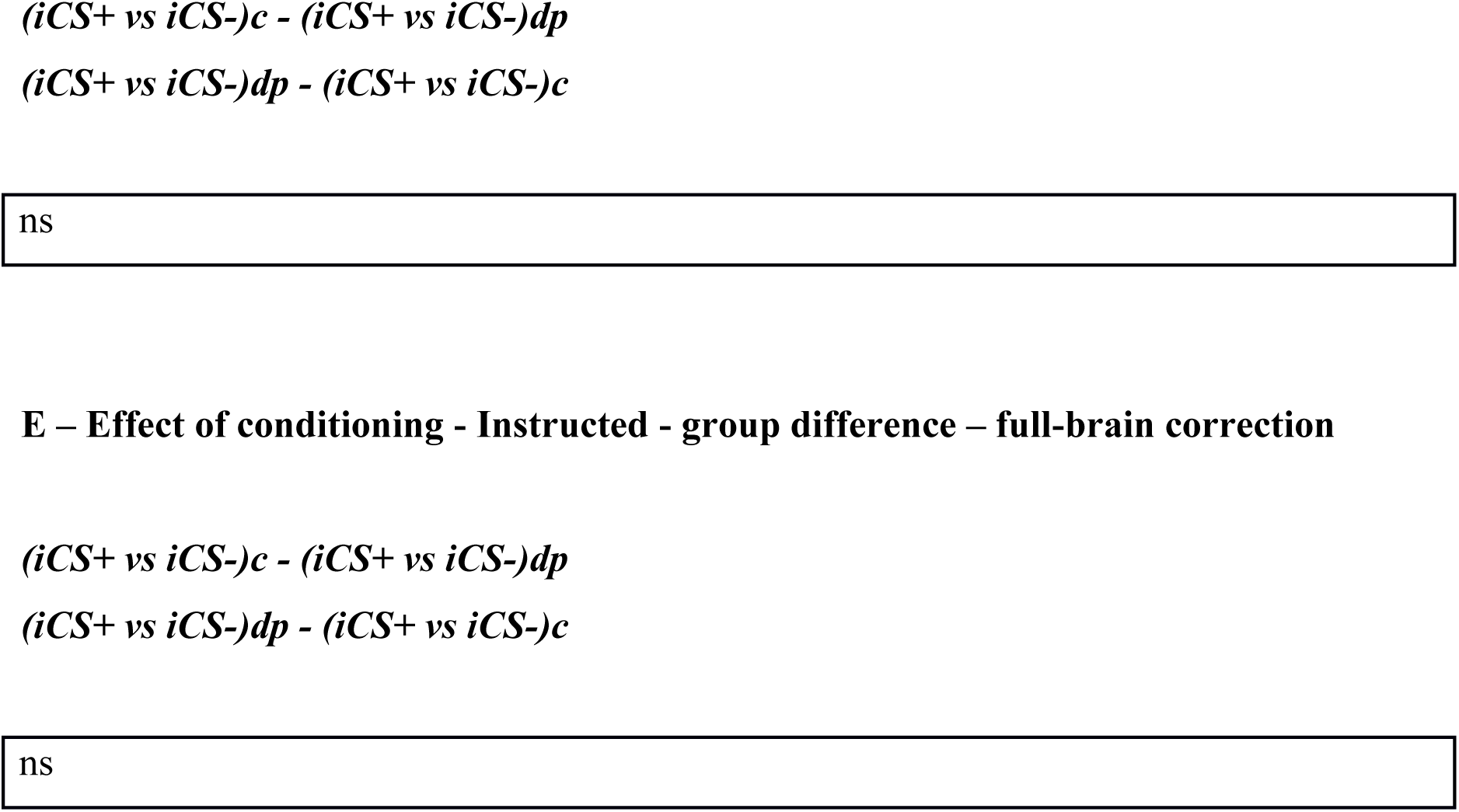
Instructed conditioning - full brain analysis. Instructed conditioning resulted in the activation of brain regions similar to the ones activated in the main effect of general conditioning. The map was thresholded at p < 0.001 (uncorrected), k > 20 transformed voxels. dp: delusion-prone group. c: control group. The shown *p-values were corrected for full brain volume (FWE-correction). ROI defined on the main effect of conditioning (Table 1) were used for small volume correction (SVC) on the group difference (E). cACC = caudal anterior cingulate cortex. dlPFC = dorsolateral prefrontal cortex.

**Table S4.**
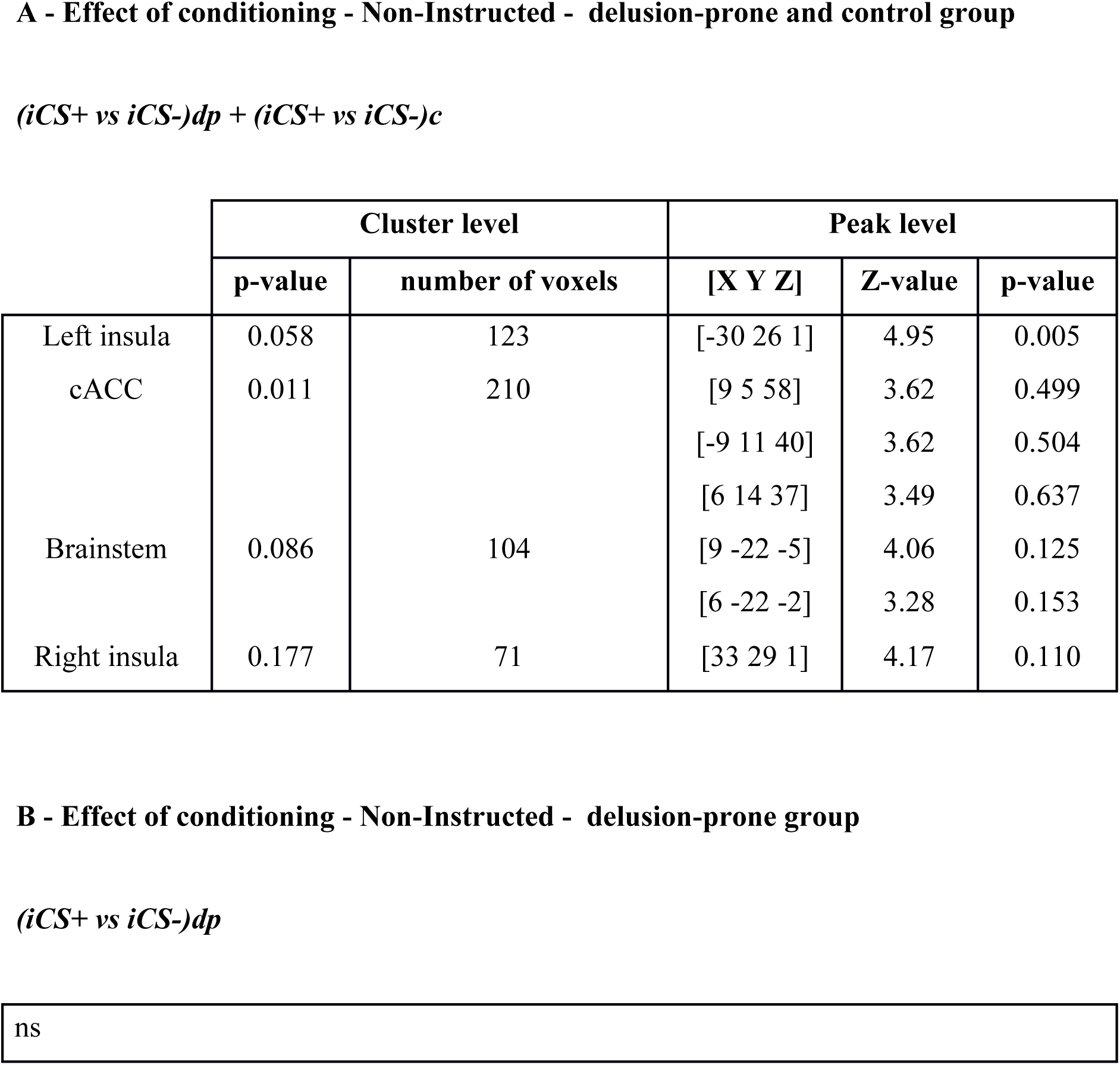
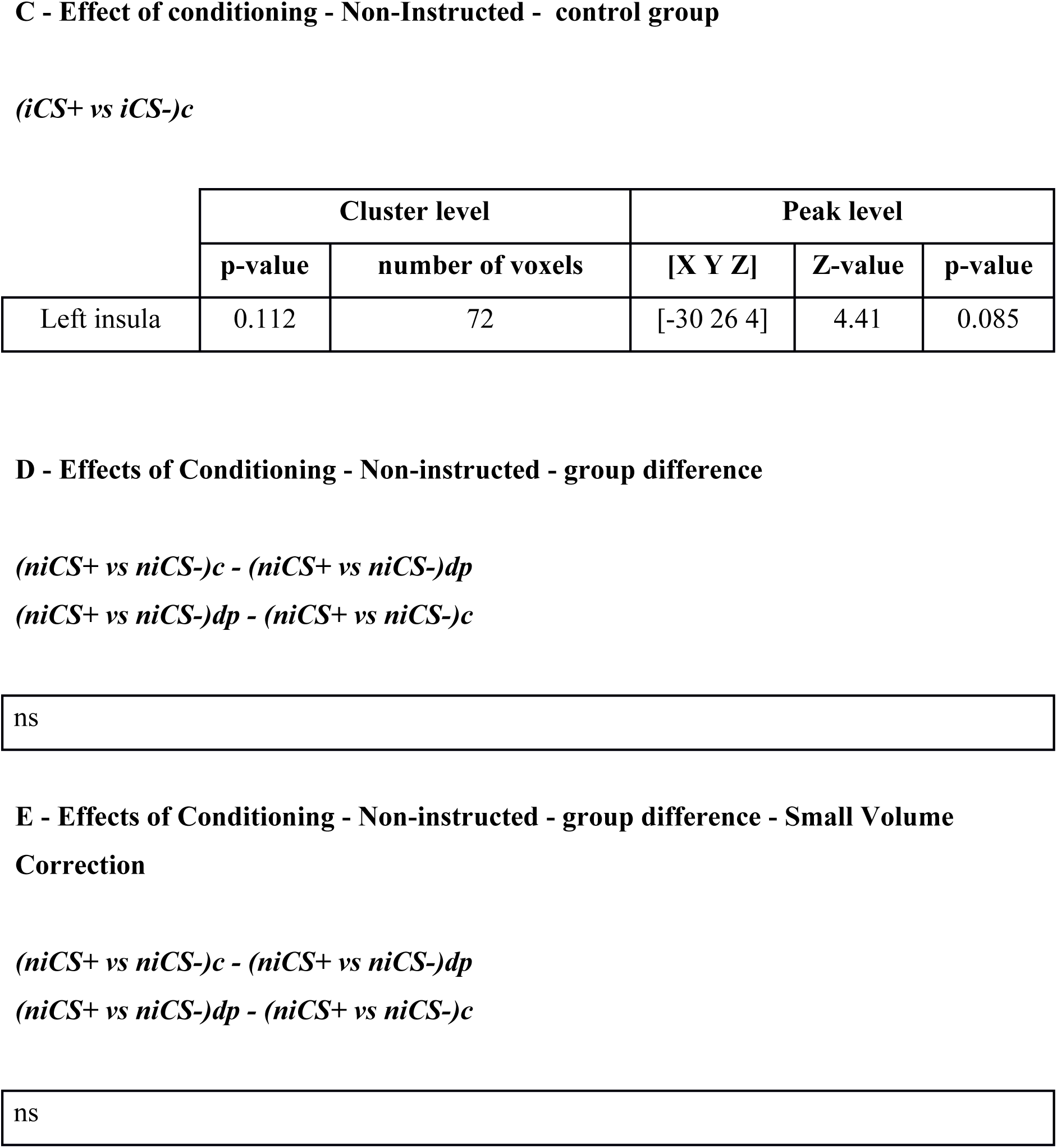
Non-Instructed conditioning - full brain analysis. Non-instructed conditioning resulted in the activation of brain regions similar to the ones activated in the main effect of general conditioning. The map was thresholded at p < 0.001 (uncorrected), k > 20 transformed voxels. dp: delusion-prone group. c: control group. The shown *p-values were corrected for full brain volume (FWE-correction). ROI defined on the main effect of conditioning (Table 1) were used for small volume correction (SVC) on the group difference (E). cACC = caudal anterior cingulate cortex.

**Table S5.** Effect of pain - full brain analysis. The main effect of pain resulted in the activation of brain regions usually found in the pain network: caudal ACC (cACC), bilateral mid- and posterior insula. We only report maximally activated voxels in cACC and posterior insula bilaterally. As there was no proper control condition for the pain, the activation pattern observed in this contrast was much wider than the one generally observed in pain studies using a low level control. The map was thresholded at p < 0.001 (uncorrected), k > 20 transformed voxels. The shown *p-values were corrected for full brain volume (FWE-correction). Peak-activations used for the subsequent PPI-analyses are shown in bold.

**Effect of pain - delusion-prone and control group**
|  | Peak level |  |  |
| --- | --- | --- | --- |
|  | Z-values | [X Y Z] | p-value* |
| cACC | Inf | [-3 2 37] | <0.0001 |
| *Right Insula* |  |  |  |
| Posterior upper Insula | Inf | **[36 -16 4]** | **<0.0001** |
| Posterior lower Insula | Inf | [39 -13 -8] | <0.0001 |
| Mid Insula | Inf | [42 11 -5] | <0.0001 |
| *Left Insula* |  |  |  |
| Mid Insula | Inf | [-39 7 -5] | <0.0001 |
| Posterior lower Insula | Inf | [-39 -10 -2] | <0.0001 |
| Posterior upper Insula | Inf | [-39 -16 13] | <0.0001 |

## References

Adams, R.A. et al., 2013. The computational anatomy of psychosis. Frontiers in psychiatry, 4(May), p.47. Available at: http://journal.frontiersin.org/article/10.3389/fpsyt.2013.00047/abstract.

Atlas, L.Y. et al., 2016. Instructed knowledge shapes feedback-driven aversive learning in striatum and orbitofrontal cortex, but not the amygdala. eLife, 5(MAY2016).

Balog, Z., Somlai, Z. & Kéri, S., 2013. Aversive conditioning, schizotypy, and affective temperament in the framework of the salience hypothesis. Personality and Individual Differences, 54(1), pp. 109–112.

Bar, M., 2003. A cortical mechanism for triggering top-down facilitation in visual object recognition. Journal of Cognitive Neuroscience, 15(4), pp. 600–609. Available at: http://www.mitpressjournals.org/doi/abs/10.1162/089892903321662976%5Cnpapers3://publication/doi/10.1162/089892903321662976.

Buchy, L., Woodward, T.S. & Liotti, M., 2007. A cognitive bias against disconfirmatory evidence (BADE) is associated with schizotypy. Schizophrenia Research, 90(1–3), pp. 334–337.

Corlett, P.R. et al., 2007. Disrupted prediction-error signal in psychosis: Evidence for an associative account of delusions. Brain, 130(9), pp. 2387–2400.

Corlett, P.R. & Fletcher, P.C., 2012. The neurobiology of schizotypy: Fronto-striatal prediction error signal correlates with delusion-like beliefs in healthy people. Neuropsychologia, 50(14), pp. 3612–3620.

Eippert, F. et al., 2007. Regulation of emotional responses elicited by threat-related stimuli. Human Brain Mapping, 28(5), pp. 409–423.

Fletcher, P.C. & Frith, C.D., 2009. Perceiving is believing: a Bayesian approach to explaining the positive symptoms of schizophrenia. Nature reviews. Neuroscience, 10(1), pp. 48–58. Available at: http://dx.doi.org/10.1038/nrn2536.

Friston, K., 2005. A theory of cortical responses. Philosophical transactions of the Royal Society of London. Series B, Biological sciences, 360(1456), pp. 815–36. Available at: http://www.pubmedcentral.nih.gov/articlerender.fcgi?artid=1569488&tool=pmcentrez&rendertype=abstract.

Friston, K.J. et al., 1997. Psychophysiological and modulatory interactions in neuroimaging. NeuroImage, 6(3), pp. 218–229. Available at: ttp://linkinghub.elsevier.com/retrieve/pii/S1053811997902913%5Cnpapers3://publication/doi/10.1006/nimg.1997.0291.

Fullana, M.A. et al., 2016. Neural signatures of human fear conditioning: an updated and extended meta-analysis of fMRI studies. Molecular Psychiatry, 21(4), pp. 500–508. Available at: http://www.nature.com/doifinder/10.1038/mp.2015.88.

Fusar-Poli, P. et al., 2013. The psychosis at risk state: a comprehensive state-of-the-art review. JAMA Psychiatry, 70(1), pp. 107–120. Available at: http://archpsyc.jamanetwork.com/article.aspx?articleid=1392281.

Golkar, A. et al., 2012. Distinct Contributions of the Dorsolateral Prefrontal and Orbitofrontal Cortex during Emotion Regulation. PLoS ONE, 7(11).

Holt, D.J. et al., 2009. Extinction Memory Is Impaired in Schizophrenia. Biological Psychiatry, 65(6), pp. 455–463.

Holt, D.J. et al., 2012. Failure of neural responses to safety cues in schizophrenia. Archives of general psychiatry, 69(9), pp. 893–903. Available at: http://archpsyc.jamanetwork.com/article.aspx?doi=10.1001/archgenpsychiatry.2011.2310%5Cnpapers3://publication/doi/10.1001/archgenpsychiatry.2011.2310.

Javitt, D.C. & Freedman, R., 2015. Sensory processing dysfunction in the personal experience and neuronal machinery of schizophrenia. American Journal of Psychiatry, 172(1), pp. 17–31.

Jensen, J. et al., 2008. The formation of abnormal associations in schizophrenia: neural and behavioral evidence. Neuropsychopharmacology : official publication of the American College of Neuropsychopharmacology, 33(3), pp. 473–9. Available at: http://www.ncbi.nlm.nih.gov/pubmed/17473838.

Kanske, P. et al., 2011. How to regulate emotion? Neural networks for reappraisal and distraction. Cerebral Cortex, 21(6), pp. 1379–1388.

Kapur, S., 2003. Psychosis as a state of aberrant salience: A framework linking biology, phenomenology, and pharmacology in schizophrenia. American Journal of Psychiatry, 160(1), pp. 13–23.

Kuepper, R., Skinbjerg, M. & Abi-Dargham, A., 2012. The dopamine dysfunction in schizophrenia revisited: New insights into topography and course. Handbook of Experimental Pharmacology, 212, pp. 1–26.

Louzolo, A. et al., 2017. Delusion-proneness displays comorbidity with traits of autistic-spectrum disorders and ADHD. PloS one, 12(5), p.e0177820.

McLean, B.F., Mattiske, J.K. & Balzan, R.P., 2016. Association of the jumping to conclusions and evidence integration biases with delusions in psychosis: A detailed meta-analytic approach. Schizophrenia Bulletin, pp. 1–11.

Mertens, G. et al., 2016. Fear expression and return of fear following threat instruction with or without direct contingency experience. Cognition and Emotion, 30(5), pp. 968–984.

Moritz, S. & Woodward, T.S., 2006. A generalized bias against disconfirmatory evidence in schizophrenia. Psychiatry Research, 142(2–3), pp. 157–165.

Murray, G.K. et al., 2008. Substantia nigra/ventral tegmental reward prediction error disruption in psychosis. Molecular psychiatry, 13(3), pp. 239, 267–76. Available at: http://dx.doi.org/10.1038/sj.mp.4002058.

Orenes, I. et al., 2012. Schizotypal people stick longer to their first choices. Psychiatry Research, 200(2–3), pp. 620–628.

van Os, J. et al., 2009. A systematic review and meta-analysis of the psychosis continuum: evidence for a psychosis proneness–persistence–impairment model of psychotic disorder. Psychological Medicine, 39(2), p. 179. Available at: http://www.journals.cambridge.org/abstract_S0033291708003814.

Peters, E. et al., 2004. Measuring Delusional Ideation: The 21-Item Peters et al. Delusions Inventor....: Joshua. Schizophrenia Bulletin, 30(4), pp. 1005–1022. Available at: http://eds.a.ebscohost.com.libproxy.mbts.edu/eds/pdfviewer/pdfviewer?vid=6‖sid=c17f1aa0-7c94-4184-bbe4-9767a3e94354%40sessionmgr4003‖hid=4102.

Petrovic, P. et al., 2010. A prefrontal non-opioid mechanism in placebo analgesia. Pain, 150(1), pp. 59–65.

Petrovic, P. et al., 2008. Oxytocin attenuates affective evaluations of conditioned faces and amygdala activity. The Journal of neuroscience : the official journal of the Society for Neuroscience, 28(26), pp. 6607–15. Available at: http://www.pubmedcentral.nih.gov/articlerender.fcgi?artid=2647078&tool=pmcentrez&rendertype=abstract.

Petrovic, P. et al., 2002. Placebo and opioid analgesia--imaging a shared neuronal network. Science (New York, N.Y.), 295(5560), pp. 1737–40. Available at: http://www.sciencemag.org/content/295/5560/1737.full.html%5Cnhttp://www.sciencemag.org/content/295/5560/1737.full.html%5Cnhttp://www.sciencemag.org/content/suppl/2002/02/27/1067176.DC1.html%5Cnhttp://www.sciencemag.org/content/295/5560/1737.full.html#related-urls%5Cnhttp://www.sciencemag.org/cgi/collection/neurosci.

Petrovic, P. et al., 2005. Placebo in emotional processing - Induced expectations of anxiety relief activate a generalized modulatory network. Neuron, 46(6), pp. 957–969.

Petrovic, P. & Castellanos, F.X.M., 2016. Top-down dysregulation - from ADHD to emotional instability. Frontiers in Behavioral Neuroscience, 10, p. 70. Available at: http://journal.frontiersin.org/article/10.3389/fnbeh.2016.00070/abstract.

Phelps, E. et al., 2001. Activation of the left amygdala to a cognitive representation of fear. Nature Neuroscience, 4(4), pp. 437–441. Available at: http://www.nature.com/doifinder/10.1038/86110%5Cnpapers3://publication/doi/10.1038/86110.

Powers, A.R., Mathys, C. & Corlett, P.R., 2017. Pavlovian conditioning-induced hallucinations result from overweighting of perceptual priors. Science (New York, N.Y.), 357(6351), pp. 596–600.

Roiser, J.P. et al., 2009. Do patients with schizophrenia exhibit aberrant salience? Psychological Medicine, 39(2), p. 199-. Available at: http://www.journals.cambridge.org/abstract_S0033291708003863.

Romaniuk, L. et al., 2010. Midbrain activation during Pavlovian conditioning and delusional symptoms in schizophrenia. Archives of general psychiatry, 67(12), pp. 1246–1254.

Schlagenhauf, F. et al., 2014. Striatal dysfunction during reversal learning in unmedicated schizophrenia patients. NeuroImage, 89, pp. 171–180.

Schmack, K. et al., 2013. Delusions and the role of beliefs in perceptual inference. The Journal of neuroscience : the official journal of the Society for Neuroscience, 33(34), pp. 13701–12. Available at: http://www.ncbi.nlm.nih.gov/pubmed/23966692.

Schmack, K. et al., 2017. Enhanced predictive signalling in schizophrenia. Human Brain Mapping, 38(4), pp. 1767–1779.

Tanasescu, R. et al., 2016. Functional reorganisation in chronic pain and neural correlates of pain sensitisation: A coordinate based meta-analysis of 266 cutaneous pain fMRI studies. Neuroscience and Biobehavioral Reviews, 68, pp. 120–133.

Teufel, C. et al., 2010. Deficits in sensory prediction are related to delusional ideation in healthy individuals. Neuropsychologia, 48(14), pp. 4169–4172.

Teufel, C. et al., 2015. Shift toward prior knowledge confers a perceptual advantage in early psychosis and psychosis-prone healthy individuals. Proceedings of the National Academy of Sciences, 112(43), pp. 13401–13406. Available at: http://www.pnas.org/lookup/doi/10.1073/pnas.1503916112%5Cnhttp://www.pnas.org/content/early/2015/10/06/1503916112.abstract.

Veckenstedt, R. et al., 2011. Incorrigibility, jumping to conclusions, and decision threshold in schizophrenia. Cognitive neuropsychiatry, 16(2), pp. 174–192. Available at: http://www.tandfonline.com/doi/abs/10.1080/13546805.2010.536084.

Wager, T.D. et al., 2008. Prefrontal-Subcortical Pathways Mediating Successful Emotion Regulation. Neuron, 59(6), pp. 1037–1050.

Wager, T.D. & Atlas, L.Y., 2015. The neuroscience of placebo effects: Connecting context, learning and health. Nature Reviews Neuroscience, 16(7), pp. 403–418. Available at: http://www.nature.com/doifinder/10.1038/nrn3976.

Woodward, T.S. et al., 2007. A bias against disconfirmatory evidence is associated with delusion proneness in a nonclinical sample. In Schizophrenia Bulletin. pp. 1023–1028.

Woodward, T.S. et al., 2008. Belief inflexibility in schizophrenia. Cognitive Neuropsychiatry, 13(3), pp. 267–277. Available at: http://www.tandfonline.com/doi/abs/10.1080/13546800802099033.

Woodward, T.S., Moritz, S. & Chen, E.Y.H., 2006. The contribution of a cognitive bias against disconfirmatory evidence (BADE) to delusions: A study in an Asian sample with first episode schizophrenia spectrum disorders. Schizophrenia Research, 83(2–3), pp. 297–298.

